# Double-stranded DNA reduces dsRNA degradation in the saliva and significantly enhanced RNAi-mediated gene silencing in *Halyomorpha halys*

**DOI:** 10.1101/2024.10.10.617619

**Authors:** Venkata Partha Sarathi Amineni, Georg Petschenka, Aline Koch

## Abstract

The invasive pest *Halyomorpha halys* (Hemiptera: Pentatomidae) poses a significant threat to agriculture, necessitating effective control methods beyond chemical pesticides. Our research explores RNA interference (RNAi) as a targeted gene silencing approach for *H. halys* population management. However, the variable efficacy of RNAi across insect orders, particularly in hemipteran insects like *H. halys*, poses challenges. Ex vivo degradation assays revealed rapid degradation of double-stranded RNA (dsRNA) in *H. halys* (*Hh*) saliva and extracts of salivary glands across several growth stages, attributed to the high expression of the DNA/RNA non-specific nuclease (*HhNSE)*. A key discovery from our research was that double-stranded DNA (dsDNA) can act as a protective agent, increasing the stability of dsRNA in saliva probably by competitive inhibition of *HhNSE*. Based on the well-established lethality of silencing the gene encoding the heavy chain of clathrin (*HhCHC*) in insects, this gene was chosen as a target to test the functionality of our dsDNA-based formulation for enhancing dsRNA-mediated gene silencing. Our in-vivo experiments showed increased *HhCHC* silencing after 72 hours of feeding initiation with a mixture of dsRNA-CHC and dsDNA, as opposed to dsRNA alone. This discovery indicates potential for enhancing the efficiency of orally delivered dsRNA through formulations based on dsDNA. Although the injection of dsRNA-CHC resulted in near-total mortality, the dsDNA formulation did not significantly enhance mortality rates when fed together with dsRNA-CHC. This emphasises the necessity for further investigation into additional factors beyond nuclease activity, such as the understanding of dsRNA uptake and release mechanisms within the gut epithelial cells of *H. halys*. Nevertheless, our study opens avenues for developing cost-effective formulations to enhance RNAi efficacy in *H. halys* and perhaps other insects where nucleases hinder dsRNA delivery, representing a promising solution for sustainable pest control.

**Graphical Abstract:** 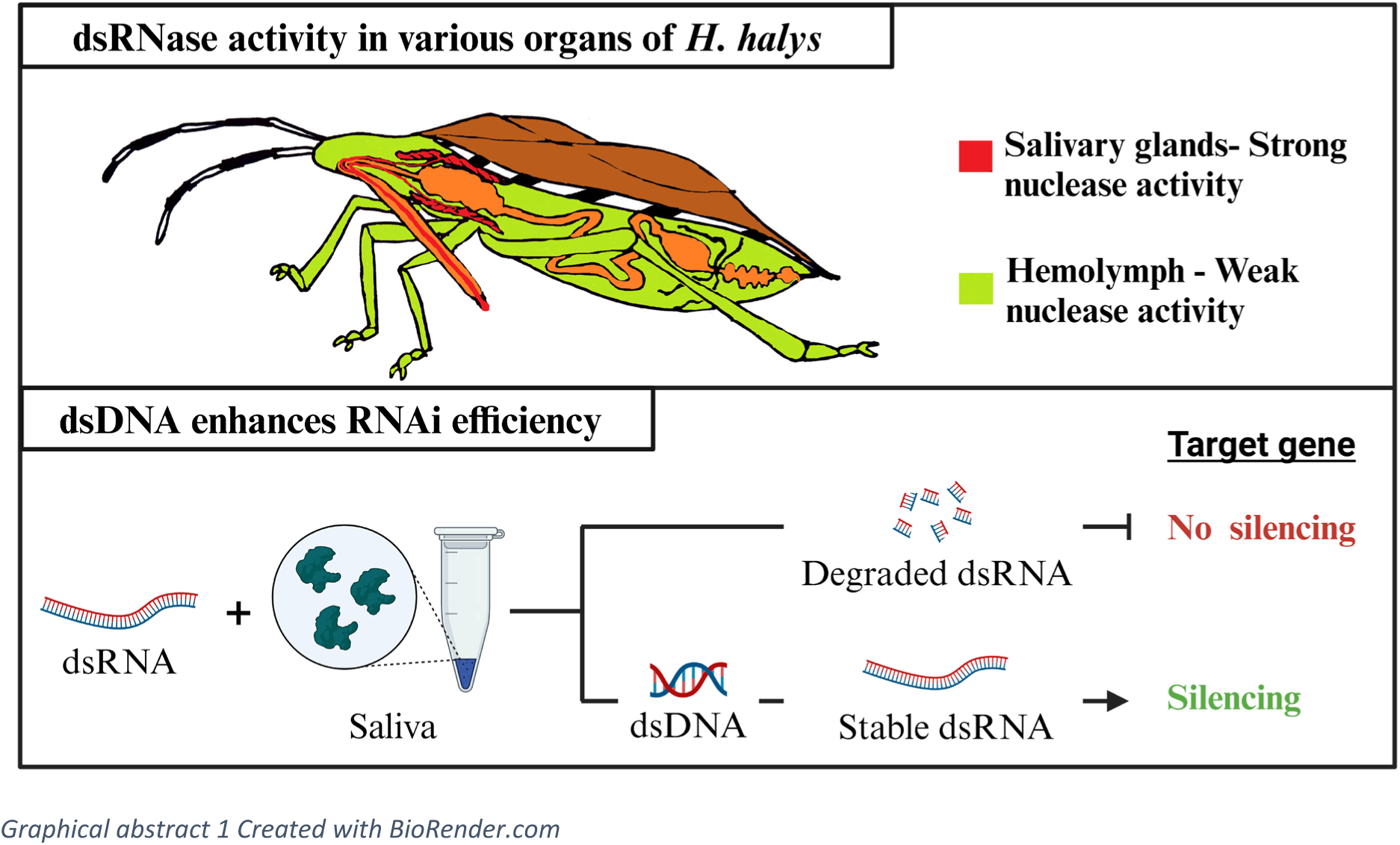

## 1. Introduction

The global demand for effective, species-specific (i.e. selective), and sustainable alternatives to conventional chemical pesticides in agriculture is growing^1^. RNA interference (RNAi) is a mechanism of gene regulation where short, non-coding RNA molecules suppress the expression of complementary target genes (gene silencing). RNAi evolved as a natural defense against viruses in plants^2^, mammals^3,4^, and insects^5^. Modern RNAi-based plant protection technologies utilize molecular mechanisms of post-transcriptional gene silencing, which are initiated by Dicer, an RNase III-like enzyme, breaking down a long double-stranded (ds) RNA precursor into 21 to 24 nucleotide (nt) small interfering (si)RNA duplexes^6^. Post-transcriptional gene silencing occurs in the cytoplasm of the cell, where the siRNAs are incorporated into an RNA-induced silencing complex (RISC)^7^ that contains an Argonaute protein with an RNA-binding domain and endonucleolytic activity. In an ATP-dependent reaction, siRNA is separated by RISC, resulting in a sense and an antisense strand. The sense strand, which is identical to the cell’s endogenous mRNA, is degraded while the antisense strand remains bound to the RISC and targets complementary mRNA transcripts for degradation^8^.

The efficiency of RNAi-based control of insect pests varies drastically among insect orders (for an overview, see^9^). For example, 100% mortality were reached in the inbred German laboratory D01 strain of *Leptinotarsa decemlineata* (Coleoptera; Chrysomelidae) when fed with 30 ng dsRNA-actin per single 2^nd^ instar larva on a 2 cm diameter leaf disc^10^. In contrast, Lepidopterans such as *Spodoptera exigua*^11^ and *Bombyx mori*^12^, and Hemipterans, such as *Plautia stali*^13^, *Euschistus heros*^14^, and *Acyrthosiphon pisum*^15^, were less sensitive to dsRNA when treated with relatively higher quantities of dsRNA per individual via feeding. Numerous factors likely contribute to this strong variation of RNAi efficiency including the degradation of dsRNA by nucleases in the saliva and gut of *Nezara viridula*^16^ and the hemolymph of *Spodoptera frugiperda*^17^. Additionally, one of the major impediments to RNAi efficiency in *S. frugiperda* is the accumulation of dsRNA in endosomes in cultured (Sf9), midgut and fat body cells^18^. To bypass the challenge of dsRNA degradation by nucleases and other physiological barriers, protective formulations need to be developed which facilitate the delivery of dsRNA into insect target insect cells for inducing a proper post-transcriptional gene silencing response.

*Halyomorpha halys* (Stål, 1855) (Hemiptera, Pentatomidae), the brown marmorated stink bug, is a pest invasive to Europe and the Americas. *H. halys* is a hemimetabolous insect that goes through five nymphal stages before becoming an adult^19^. Due to its wide host range and high reproductive output, it is responsible for severe economic losses in agriculture^20^. For example, *H. halys* was first reported in 2012 in Italy^21^ and resulted in over EUR 588 million damage in the year 2019^22^. To date, broad-range chemical insecticides such as carbamates, neonicotinoids and pyrethroids etc., are the only effective control method for *H. halys* but pose environmental risks^23,24^. Due to its high potential for selectivity, RNAi represents a promising method for sustainable *H. halys* management^25–27^. However, high nuclease activity in the saliva and midgut extracts of adult *H. halys*^27,28^ might limit the delivery of dsRNA and thus, the efficiency of target gene silencing.

The DNA/RNA non-specific nucleases (NSE) show a broad substrate affinity and digest dsRNA, dsDNA, single stranded (ss)RNA, ssDNA, and RNA/DNA hybrids^29,30^. Because NSEs degrade full length dsRNA efficiently, they are also referred to as dsRNases^31^. An NSE was first identified in the midgut homogenate and liquid phase of the midgut of *B. mori* larvae^12,32^, and later studies revealed the strong expression of *NSE*s in a variety of insects^17,33–35^ degrading dsRNA in oral cavity, gut lumen and hemolymph before reaching the target cells. This ultimately reduces the efficiency of target gene silencing.

The clathrin protein plays a crucial in the uptake of nutrients through the endocytosis pathway in eukaryotic cells^36,37^. Triskelion structured clathrin consists of three 190 kDa heavy chains and a single light chain (∼25 kDa), which connect at their C-terminal ends to constitute the fundamental clathrin subunit^38^. Knocking down the *clathrin heavy chain* (*CHC*) gene has been shown to result in significant mortality across various insect groups^39,40^.

In this study, we aimed to evaluate a potential solution for protecting dsRNA from nuclease degradation when administered orally, which could further optimize the efficiency of RNAi in *H. halys*. First, we evaluated the stability of dsRNA in saliva, salivary gland extract and hemolymph of *H. halys* across different developmental instars. To determine whether dsRNA degradation could be mediated by nucleases, we quantified the relative expression of ribonucleases genes, specifically DNA/RNA *non-specific nuclease* (*HhNSE*), *exoribonuclease-1* (*HhEri-1*), and *small RNA degrading nuclease-1* (*HhSDN-1*), in the salivary glands of *H. halys* using RT-PCR. Furthermore, we explored the capacity of both PCR amplified and ultrapure salmon-sperm dsDNAs to act as a competitive inhibitor of HhNSEs and subsequently reduce dsRNA degradation in *H. halys* saliva through an ex-vivo dsRNA degradation assay. Finally, we performed in vivo assays by orally administering dsRNA-CHC targeting *HhCHC* (XM_014431604.1), combined with dsDNA, to examine the efficacy of this formulation in enhancing oral RNAi efficiency. Our findings suggest that formulating dsRNA with dsDNA can bypass the severe dsRNase activity in the saliva, thereby enhancing oral RNAi efficiency in *H. halys*.

## 2. Materials and methods

### 2.1 Insect rearing

*H. halys* was obtained from the laboratory of Katz Biotech AG, Germany, and reared on a diet composed of fresh green beans, carrots, and husked sunflower seeds. Insects were supplied with water in 15 ml falcon tubes plugged with cotton wicks. We used the same type of aerated plastic containers (19x19x19 cm) for all developmental stages which were lined with kitchen paper and maintained in Fitotron HGC1014 Modular Growth Chambers (Weiss Technik GmbH, Reiskirchen, Germany) at 25 °C, 65% humidity, and a photoperiod of 16:8 hours Insects were provided with fresh food three times per week and transferred to new boxes once per week.

### 2.2 dsRNA and dsDNA acquisition

For the ex-vivo dsRNA degradation studies, beta-glucuronidase dsRNA (dsRNA-GUS) of 225 base pairs (bp) was purchased from RNAGreentech LLC (Frisco, USA). dsRNA-CHC and dsRNA-GFP used in the in-vivo assays was acquired from Genolution AgroRNA (Seoul, Korea), see dsRNA sequences in Table. S1. Two different dsDNAs were used in this study: a 246 bp purified PCR product from *Arabidopsis thaliana, Actin-2* (*dsDNA-AtACT*, accession number NM_001338359.1) and another 102 bp fragment of *H. halys Argonaute-2 like protein* (*dsDNA-HhAGO2*, accession number XM_024358503.1). The degradation experiments were repeated using shorter *dsDNA-HhAGO2* fragments, because the dsRNA-GUS fragment became difficult to visualise on the agarose gel when the smear of the degraded *dsDNA-AtACT* fragment overlapped with it. The primers used are listed in supplementary table 1. PCR was performed using ALLinTM HiFi DNA Polymerase (highQu GmbH, Kraichtal, Germany), and the amplified dsDNA during the PCR reaction was purified with Wizard SV Gel and a PCR Clean-Up System (Promega). Subsequently, the quality of dsDNAs was assessed under UV light by gel electrophoresis at 100 V for 30 minutes on an in-gel stained 1% agarose gel with Midori Advance Green (NIPPON Genetics EUROPE, Düren, Germany) in 1x TAE buffer.

### 2.3 Saliva, salivary gland extract, and hemolymph isolation

Collection of saliva was restricted to fourth and fifth instar nymphs and adults, as described by^41^ Peiffer and Felton (2014), due to the practical difficulties of collecting from younger stages. During sample collection, insects were placed on ice for five minutes, then turned on their backs and fixed on double-sided adhesive tape under a stereomicroscope to prevent movement during saliva collection. After 1-2 minutes at room temperature (ca. 22-25 °C), bugs released several droplets of saliva at the tip of their proboscis which were collected using nuclease-free 10 µL pipette tips and pooled in Eppendorf tubes on ice to reach an approximate amount of 0.2 to 1 µL from each adult, 0.2-0.3 µL from each 5^th^ instar, and 0.1-0.2 µL from each 4^th^ instar insect. Samples were stored on ice until testing.

Salivary glands (including accessory gland, posterior and anterior lobes) from all instars except instar 1 were gently removed from the body by pulling out the head horizontally under ice-cold PBS (pH 7.4) to minimize cell damage under the stereomicroscope. The glands were separated from the head, rinsed for five seconds in fresh PBS, and transferred to a 1.5 ml tube on ice. Between seven and ten intact salivary glands per developmental stage were pooled and kept on ice. Next, 3 to 4 µL of PBS were added to the glands and tubes were slightly vortexed for 10 seconds and centrifuged at 1,000 x g for 15 minutes at 4 °C. The resulting supernatant was collected and used for degradation assays. The procedure for extracting salivary gland extract was partially adapted from Fischer et al 2020 ^42^. We thank Dr. Maike Fischer and Dr. Heiko Vogel for their valuable advice on the extraction of salivary gland contents, which contributed to the refinement of our protocol.

Hemolymph was collected from adults and all developmental stages except first and second instar nymphs due to their small body size. An incision was made at the distal portion of the intersegmental membrane between femur and tibia using sterilized spring scissors, and the hemolymph collected with nuclease-free 10 µL pipette tips was stored in an ice-cold 1.5 ml tube containing a few crystals (about the size of a sesame seed) of N-phenylthiourea (Sigma-Aldrich, Germany) to prevent melanization^43^. Hemocytes were removed by centrifugation at 1,000 x g for 8 min at 4 °C^44^, and the supernatant was used for the degradation assays. Degradation assays were performed at least in duplicate for all sample types (i.e., saliva, salivary gland contents, and hemolymph) except for the fourth and fifth stages of saliva (supplementary data section 4).

### 2.4 *Ex vivo* dsRNA stability assay

For all ex vivo dsRNA degradation assays, 2 µL of each of the three freshly collected samples (saliva, salivary gland extracts, and hemolymph) were incubated undiluted with 20 µL of GUS-dsRNA (100 ng/µL) at 25 °C using a 1.5 ml Thermomixer (Eppendorf). At ∼1, 10, 30, 60, 120, and 240 minutes after incubation, aliquots of 5 µL were collected and combined with 2 µL of 1% SDS (sodium dodecyl sulfate) in water to stop the nuclease reaction^34^. The samples were immediately subjected to gel electrophoresis analysis after the experiment. Using 1x TAE buffer, an 1% agarose gel was prepared, in-gel stained with Midori Advance Green (NIPPON Genetics EUROPE, Düren, Germany), and electrophoresed at 100 V for 30 minutes, followed by visualisation of dsRNA under UV light.

The volume of dsRNA bands from agarose gel images was quantified using ImageJ 1.53t software^45^ and statistically analysed across incubation times using SAS version 9.4 (SAS Institute Inc., Cary, NC, USA), with subsequent data visualization conducted in JMP (ver. 17.2.0; SAS Institute, Cary, NC). The quantified gel image data was modelled using PROC MIXED from SAS 9.4, i.e., mixed modelling and the associated analysis results and graphs are provided in supplementary materials (section 5 & ex_vivo_data.pdf). Although n=2 in our data, having multiple data points at different times made it possible to model and analyse the data. All the gel images were processed and labelled using the online platform Biorender.com.

### 2.5 RT-PCR

Gene expression levels of three different dsRNA degrading nucleases (*HhNSE* (XM_024362815.1)*, HhEri-1* (XM_024361541.1)*, HhSDN-1* (XM_014423854.2)) in salivary glands were assessed. To reduce inter-individual variation, 18-20 salivary glands from ten untreated *H. halys* adults were pooled in 400 µL of a DNA/RNA-protecting solution (Monarch Total RNA Miniprep Kit, New England Biolabs Inc., Germany) for each biological replicate. Total RNA from the homogenized glands was extracted according to the manufacturer’s recommendations (Monarch Total RNA Miniprep Kit, New England Biolabs Inc., Frankfurt, Germany). The purified RNA was quantified using a BioPhotometer Plus with a μCuvette G1.0 (Eppendorf). Three biological replicates were evaluated in total.

cDNA synthesis was performed using qScript cDNA Synthesis kit (VWR International GmbH, Darmstadt, Germany). Primers for RT-PCR were designed using the PrimeQuest tool (www.idtdna.com), followed by an off-target prediction check using PrimerBlast (National Center for Biotechnology Information, Bethesda, MD, USA). The primers used in this study are listed in supplementary table 1. RT-PCR analysis was conducted with the SYBR Green JumpStart Taq ReadyMix (Sigma-Aldrich, Germany) using Bio-Rad CFX96 PCR detection system. The RT-PCR experiments were conducted with each biological replicate consisting of two technical replicates to ensure robust and reliable results. After the initial activation step at 94 °C for 2 min, 40 cycles (94°C for 15 sec, 60°C for 60 sec, and 72°C for 30 sec) were performed. RT-PCR data analysis was conducted through the Pfaffl method^46^, which considers primer efficiencies. Average Ct values of *HhEri-1* were utilized as a calibrator gene for determining the delta Ct values and calculating the ratio of relative gene expression. For data visualization and statistical analysis, including 2-fold logarithmic transformation, ANOVA and multiple comparisons with Tukey-HSD as a post hoc test, JMP (ver. 17.2.0; SAS Institute, Cary, NC) was used.

### 2.6 dsDNA competitor assay

Given that NSE has a broad affinity for nucleic acids, we speculated that dsDNA may function as a competitor to protect dsRNA from dsRNases in *H. halys* saliva. To test this hypothesis, we amplified dsDNA originating from *A. thaliana* and *H. halys* (see section 2.2). At 25 ^°^C, 2 µL of fresh, undiluted saliva was incubated with 20 µL of 100 ng/µL dsRNA-GUS and 100 ng/µL of dsDNA, together diluted in ultrapure, nuclease-free water (Invitrogen by Thermo Fisher Scientific, Waltham, MA), and aliquots were collected at ∼1, 10, 30, and 60 minutes. Before incubation, the reaction solution was vortexed and spun for 30 seconds to ensure proper mixing and to collect the solution at the bottom of the reaction tube after saliva was added. Additionally, 1:0.5, 1:2, and 1:3 ratios of dsRNA-GUS:dsDNA-AtACT were tested. At each time point, 5 µL aliquots were collected and 2 µL of 1% SDS was added to stop the nuclease reaction and left at room temperature. Upon completion of collection of all samples, samples were immediately investigated for dsRNA-GUS stability by gel electrophoresis (see section 2.2) without freezing.

### 2.7 In Vivo Assays to Assess the Efficiency of the dsRNA Formulation

This study was aimed at evaluating the effect of a dsDNA formulation on oral RNAi efficacy in second instar *H. halys*. Due to the need for large quantities for feeding bioassays, sheared UltraPure™ Salmon Sperm DNA (dsDNA-S) from ThermoFisher Scientific, Germany (catalog number: 15632011) was utilized as a formulant. The inhibitory effect of dsDNA-S against *H. halys* salivary nucleases was evaluated in line with the methodology described in section 2.6.

In conducting the bioassays, the nucleic acids of 200 µL (i.e. 1:1 ratio of dsDNA-S and dsRNA-CHC) were combined with a 1% sucrose solution and red food dye (Dr. Oetker, Germany) and packaged in stretched parafilm sachets. These sachets were immersed in fresh bush bean juice to facilitate recognition by the nymphs, then dried before being offered to the nymphs. The addition of sucrose ensured that the nymphs received the necessary carbohydrates during the incubation period.

We developed a novel method to precisely identify the feeding sites of *H. halys* nymphs on parafilm sachets (Fig. S1) as a putative indicator of feeding activity. Typically, before using their stylets to ingest sap, *H. halys* and several other stink bugs secrete a protein-rich salivary coat that coats the puncture made by their mouthparts on the host^47^. We have stained this protein sheath with the aforementioned food dye, and this method makes the coloured spots visible under the microscope and ensures that insects are feeding on the sachet.

We employed four distinct treatments: (1) feeding with 20 µg of dsRNA-GFP as a negative control; (2) 20 µg dsRNA-CHC targeting *CHC* gene in *H. halys*; (3) a combination of 20 µg dsRNA-CHC and 20 µg salmon sperm DNA (dsDNA-S) (i.e. a 1:1 ratio); and (4) direct injection of 100 ng dsRNA-CHC per nymph as a positive control. Each oral formulation was administered in 200 µL per sachet, fed to groups of five second-instar nymphs over 72 hours in a small Petri dish. Direct dsRNA injections were performed using a Beveled, 35G (NF35BV-2) needle and an UMP3 ULTRAMICROPUMP microinjector from WPI, USA, all under a stereo microscope. Following the injections, the nymphs were then allowed to feed on fresh bush beans for a duration of 72 hours (Fig. S2). Total RNA was extracted from pooled samples of 5 whole insects per biological replicate, converted to cDNA, and analysed via RT-PCR as detailed in section 2.5, with 3-9 biological replicates per treatment. mRNA levels were normalised to 18S rRNA ^27^, which served as a reference gene, and analysed using the Pfaffl method ^46^. Means of normalised log2 fold-change values between treatments were compared using one-way ANOVA followed by Tukey-HSD test.

### 2.8 dsRNA injection for survival analysis

In addition to RT-PCR assays, we carried out injection assays to assess the mortality of dsRNA-CHC. Second-instar nymphs were injected as described above. In total, we carried out three experiments with four treatments including dsRNA-CHC, dsRNA-GFP, nuclease-free (NF) water, and an untreated control. Sample sizes ranged from 10 to 37 individuals per treatment and experimental conditions were the same as described for rearing of the insects. Mortality was assessed over period of 13 days and cumulative mortality at the end of the trial was analysed by generalised linear model (GLM) using PROC GLIMMIX in SAS 9.4 (SAS Institute, Cary, NC) followed by a Tukey-HSD post-hoc test. Mortality means per treatment across all experiments were used as a main effect and ‘experiment’ was included as a covariate.

### 2.9 dsRNA and dsDNA feeding assay for survival analysis

To assess the impact of adding dsDNA to the dsRNA-CHC on the survival of *H. halys* upon oral delivery (i.e., feeding) we selected second-stage insects and offered 200 µL (100 ng/µL) of dsRNA to a group of five insects each following the procedure described for the RT-PCR experiments. Following a 72-hour feeding period, the surviving insects were transferred to a new plastic jar (dimensions of 7.5 cm in height and 8.5 cm in diameter) with a mesh-covered lid and supplied with fresh green bush beans. In this experiment, seven replicates were conducted, each comprising a specific set of treatments. Replicates 1, 2, 3, 5, and 6 comprised all seven treatments, namely dsRNA-CHC, dsRNA-CHC plus dsDNA-S (1:1), dsRNA-GFP, dsRNA-GFP plus dsDNA-S (1:1), dsDNA-S, 1% sucrose, and an untreated control. Replicate 4 comprised five treatments, with the exclusion of dsRNA-CHC plus dsDNA-S and dsRNA-GFP. Similarly, replicate 7 comprised five treatments, with the exclusion of dsRNA-CHC plus dsDNA-S, and included only dsDNA-S, 1% sucrose, and the untreated control. To confirm that the insects were feeding on the solution provided, we checked for the presence of stained feeding marks on the parafilm sachet (note that in this experiment the food dye was directly added into the feeding solution, as mentioned in section 2.7). Survival data were then recorded over a 14-day period. The mortality data were ultimately subjected to visualisation and analysis via the GLM method in JMP (version 17.2.0; SAS Institute, Cary, NC).

## 3. Results

### 3.1 Ex-vivo dsRNA stability in saliva, salivary gland extract, and hemolymph of *H. halys*

#### 3.1.1 Saliva

Incubation of dsRNA-GUS with undiluted *H. halys* saliva resulted in severe degradation of dsRNA-GUS (Fig. 1). The degree of dsRNA degradation was similar for the fourth, fifth, and adult stages of *H. halys* (Fig. 1). After one minute of incubation with saliva collected from each of the three stages, a diffuse band with a large amount of smear was already detected on the gel. After ten minutes, the dsRNA-GUS signal disappeared from the gel, indicating that the dsRNA was completely degraded.

**Figure 1.**
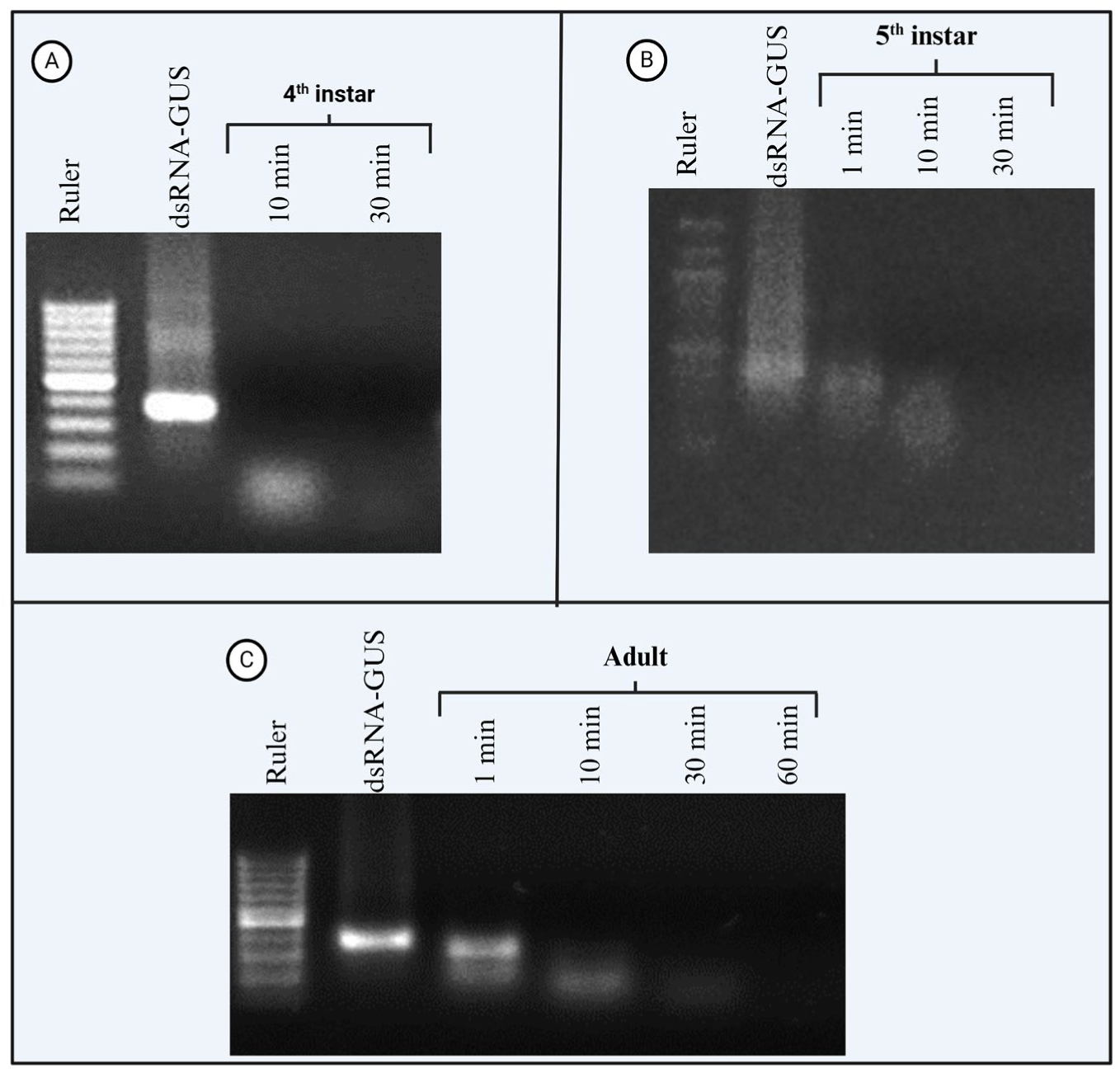
Ex vivo dsRNA degradation assay of dsRNA-GUS incubated with the saliva of *H. halys*. Saliva (2 µL) was collected from *H. halys* 4^th^ instar nymphs (A), 5^th^ instar nymphs (B), and adults incubated with 2 µg of dsRNA-GUS at 25 °C for up to 30-60 minutes, and 1% sodium dodecyl sulfate (SDS) was added to the samples to stop the nuclease reaction. The samples were run on a 1% agarose gel, and the experiment was replicated twice for each growth stage except for 4^th^ instar (n=1). In all gels, DNA ladder is used to cross-check the length of dsRNA strand and quality of the gel staining.

#### 3.1.2 Salivary glands

In addition to saliva collected from live insects, we analysed the dsRNA degradation rate in salivary gland extracts from all *H. halys* growth stages, except the first instar larvae. We found that 2 µL of salivary gland extract was sufficient to digest 2 µg of 100 ng/µL dsRNA-GUS in less than one minute (F_1, 78_ = 496.95, p < 0.001) (Fig. 2) when compared to dsRNA-GUS incubated with water. No significant difference was found between different time points (F_4, 78_ = 0.23, p = 0.918) as dsRNA-GUS degraded within 1 minute. Salivary gland extracts from all growth stages showed similar degradation activity (Fig. 2).

**Figure 2.**
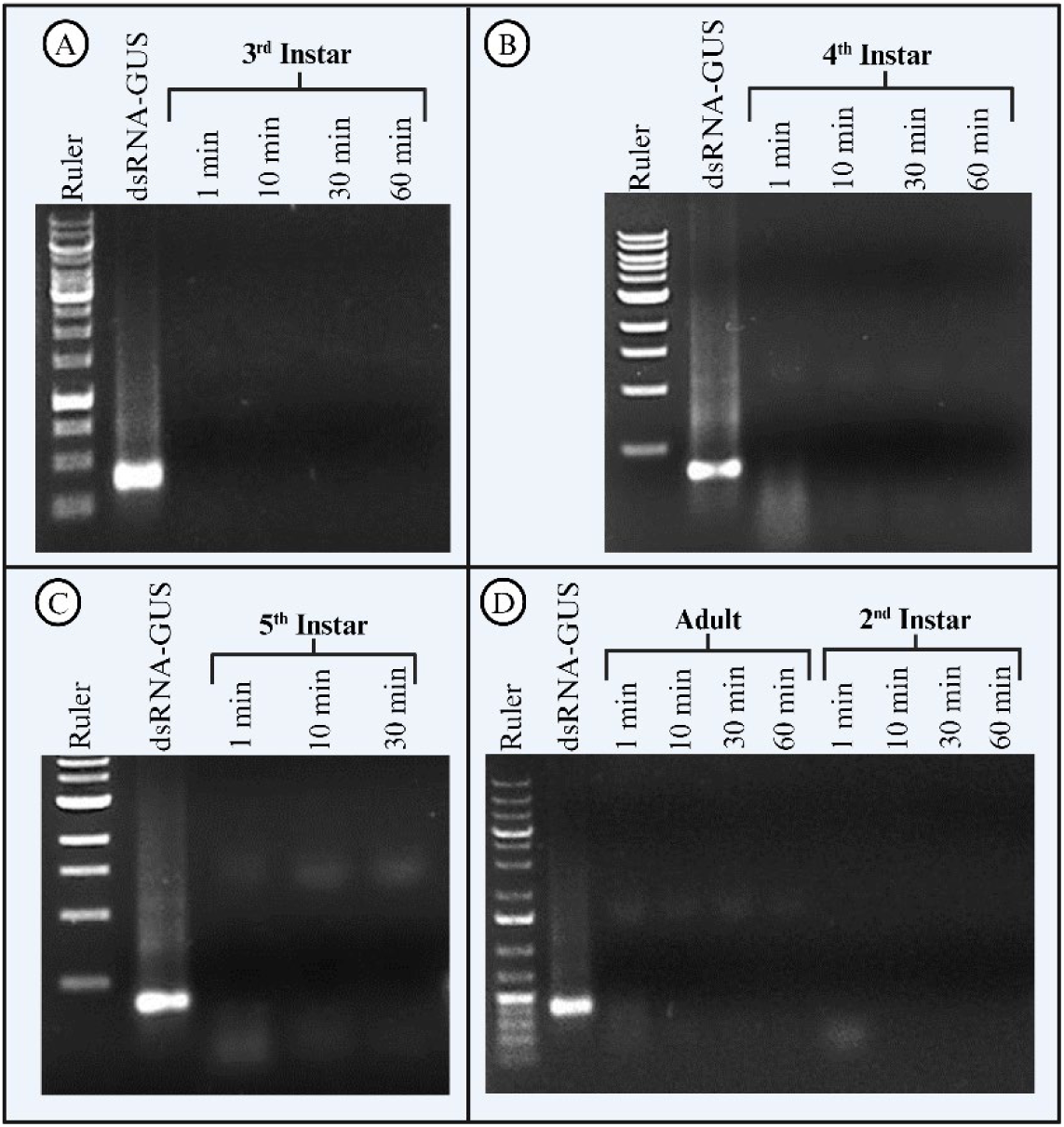
Ex vivo dsRNA degradation assay of dsRNA-GUS incubated with *H. halys* salivary gland extract. Salivary gland extract (2 µL) was extracted from *H. halys* 3^rd^ instar nymphs (A), 4^th^ instar nymphs (B), 5^th^ instar nymphs (C), and adults and 2^nd^ instar nymphs (D) and incubated with 2 µg of dsRNA-GUS for up to 60 minutes at 25 °C, and 1% SDS was added to the samples to stop the nuclease reaction. The samples were run on a 1% agarose gel and the experiment was replicated twice. DNA ladder was used in all gels.

#### 3.1.3 Hemolymph

The dsRNase activity was lower in hemolymph than in saliva or salivary gland extracts of *H. halys*, as dsRNA-GUS remained intact for up to two hours after *ex vivo* incubation (Fig. 3). Degradation of dsRNA began immediately after the start of incubation, but the degradation rate was lower compared to salivary nucleases (Fig. 1, Fig. 2). Complete degradation of dsRNA-GUS occurred earlier when dsRNA-GUS was treated with hemolymph isolated from 3^rd^ instar nymphs (Fig. 3A), where the dsRNA signal clearly started to disappear from the gel after 4 h (Fig. 3A). In contrast, the dsRNA-GUS from the 4^th^, 5^th^ instar and adult samples remained somewhat stable, while a smear in the gel indicated degradation products that were present as early as 30 min after incubation began (Fig. 3). Statistical analyses resulted that there is significant variation in dsRNA stability between different growth stages (F_1, 61.7_ = 19.24, p < 0.001) as well as between the nuclease free water and hemolymph incubated samples (F_4, 61_ = 5.54, p < 0.001).

**Figure 3.**
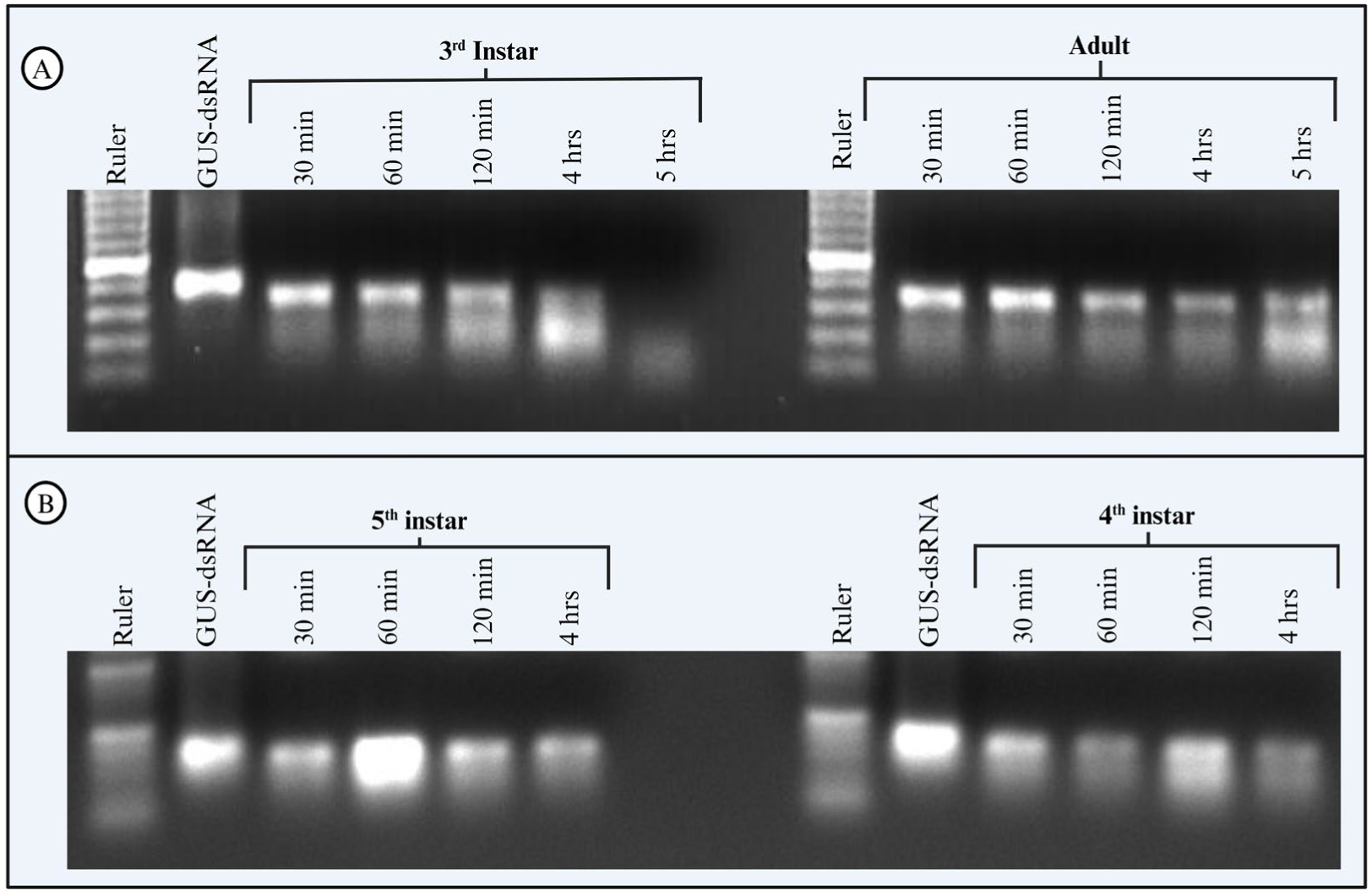
Ex-vivo dsRNA degradation assay of dsRNA-GUS incubated with *H. halys* hemolymph. Hemolymph (2 µL) from *H. halys* 3^rd^ instar nymphs and adults (A) and 5^th^ instar nymphs and 4^th^ instar nymphs (B) was incubated with 2 µg dsRNA-GUS at 25 °C for up to 4-5 hours, and 1% SDS was added to the samples to stop the nuclease reaction. The samples were run on a 1% agarose gel and the experiment was replicated twice.

### 3.2 Comparison of dsRNase expression in salivary glands

The expression levels of *HhNSE*, *HhEri-1* and *HhSDN-1* in the salivary glands of adult *H. halys* were analysed by RT-PCR, as detailed in Supplementary Table 2. It was found that *HhNSE* showed a significantly higher expression in the salivary glands compared to the other nucleases investigated, in the order *HhNSE* > *HhSDN*-1 > *HhEri-1*, as shown in figure 4. Pairwise comparisons with Tukey-HSD revealed significant differences; *HhNSE* vs. *HhEri-1* (Difference = 7.31, SE = 0.35, p < 0.001), *HhNSE* vs. *HhSDN-1* (Difference = 5.31, SE = 0.35, p < 0.001), and *HhEri-1* vs. *HhSDN-1* (Difference = -2.0, SE = 0.35, p = 0.003).

**Figure 4.**
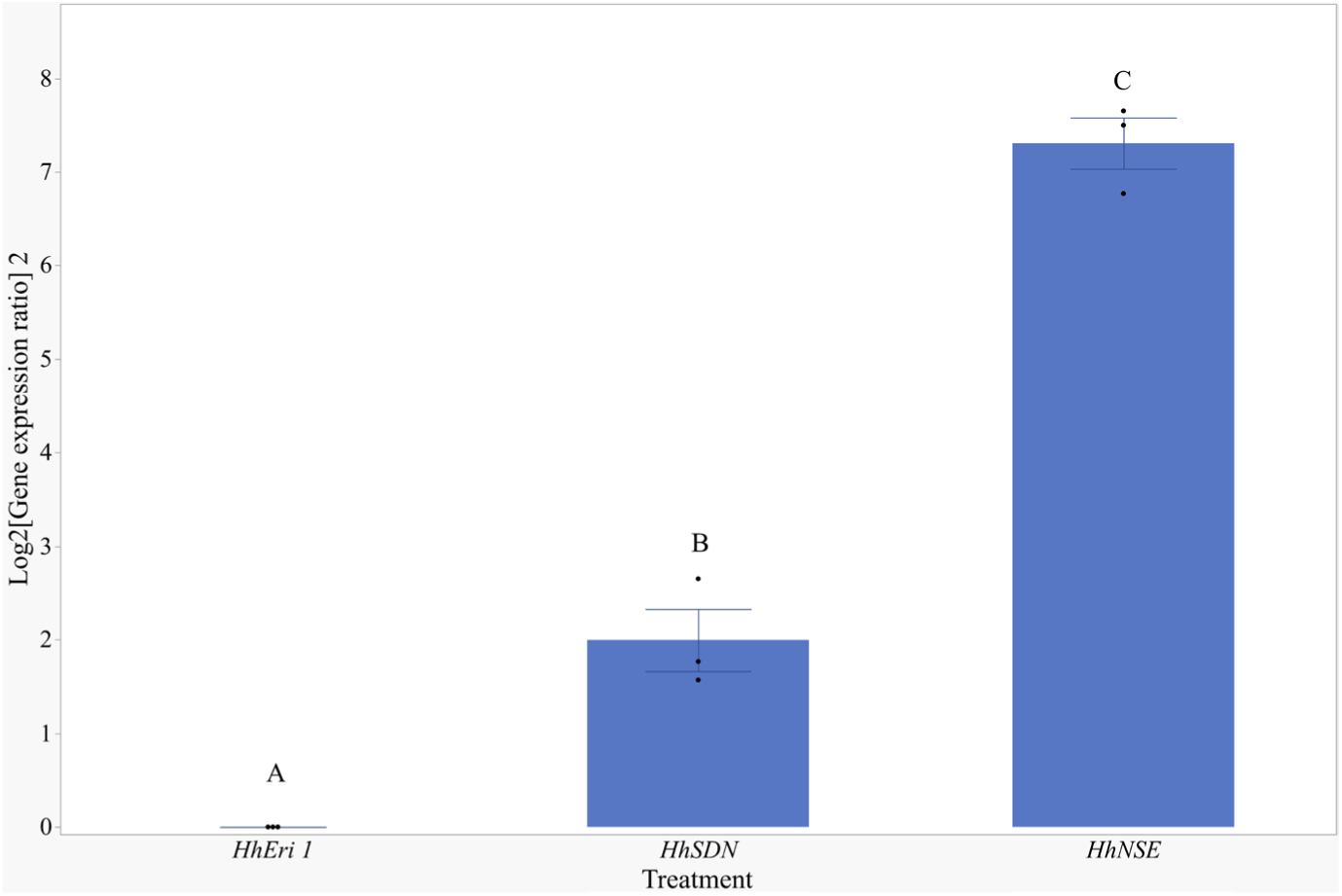
Relative expression levels of dsRNA degrading nucleases (*HhEri-1, HhSDN-1, HhNSE*) in *H. halys* salivary glands determined by RT-PCR. Total RNA from 18-20 pooled salivary glands (from ∼10 untreated *H. halys* adults) was isolated and considered as one biological sample (n). A total of three independent biological replicates (n = 3) were evaluated, with each replicate subjected to two technical replicates. The Pfaffl method was used to adjust the threshold cycle (Ct) values, taking into account primer efficiency variations, and to calculate relative gene expression ratios using *HhEri-1* as a calibrator gene. Statistical comparison of gene expression variability was conducted by ANOVA, and treatments with different letters indicate significant differences (Tukey-HSD, p < 0.01).

### 3.3 dsDNA competitively inhibits dsRNA degradation by salivary nucleases in *H. halys*

In *H. halys* salivary glands, *HhNSE* was highly expressed compared to the other known dsRNases. Therefore, *HhNSE* may be the dominant enzyme responsible for dsRNA degradation in *H. halys* saliva. The broad substrate affinity of *HhNSEs* for nucleic acids provided a potential opportunity to protect dsRNA by competitively inhibiting *HhNSEs* with dsDNA. Therefore, we investigated the stability of dsRNA when incubated with *H. halys* saliva with the addition of dsDNA. dsRNA-GUS (2 µg) was incubated with 2 µL of undiluted adult *H. halys* saliva alone and with 2 µg, 3 µg, and 4 µg of 246 bp dsDNA-*At*ACT separately. The degradation rate of dsRNA-GUS decreased with increasing concentration of 246 bp dsDNA-AtACT (Fig. 5A). A distinct band of dsRNA-GUS was detected at 30 minutes when dsRNA-GUS was co-incubated with dsDNA-AtACT and saliva samples at all dsRNA:dsDNA ratios (Fig. 5A). In contrast, the 2 µL of saliva from the same stock completely degraded the 4 µg of dsRNA-GUS within 30 minutes of incubation when no dsDNA was added (Fig. 5B).

**Figure 5.**
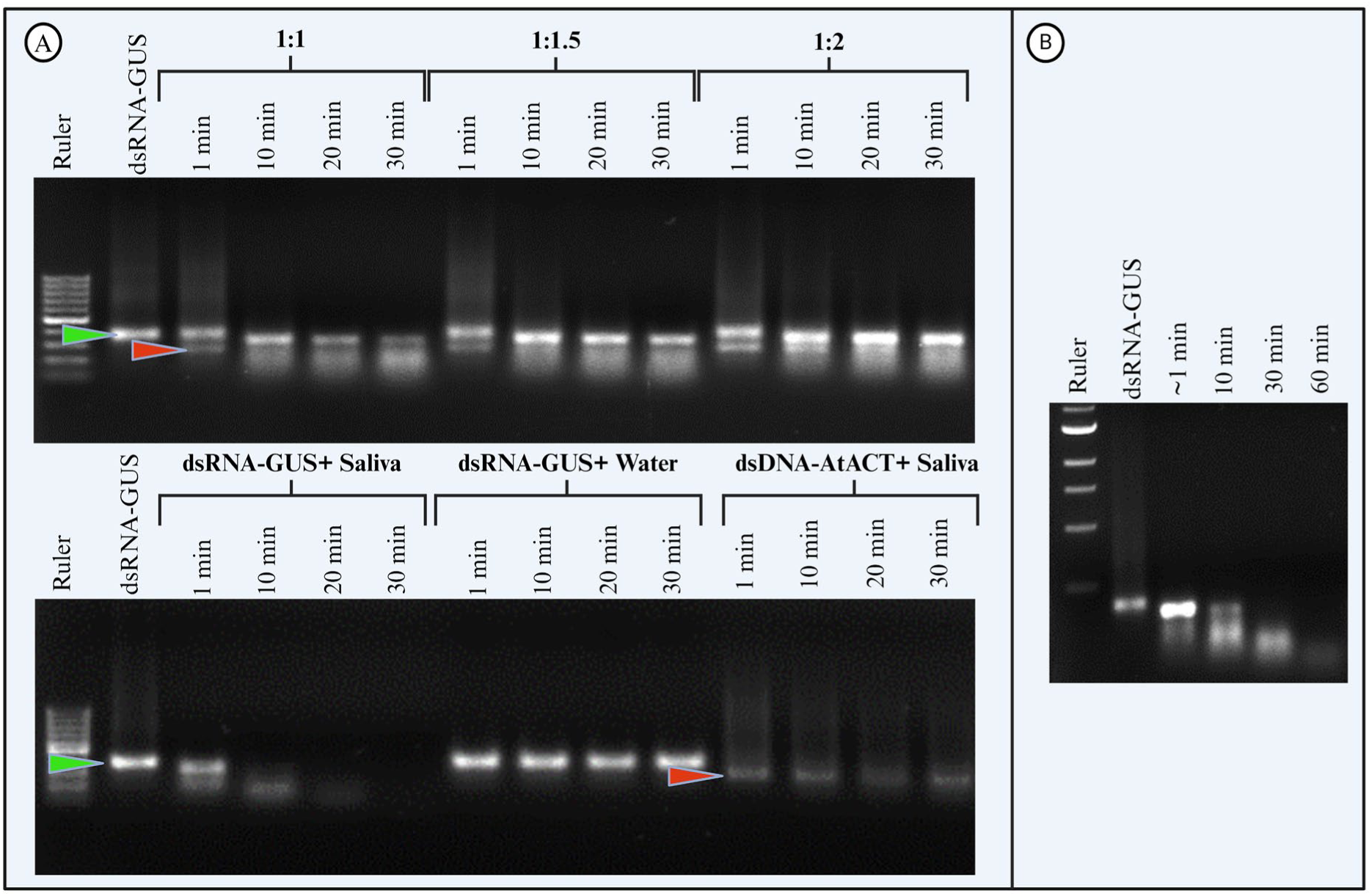
Testing the stability of dsRNA-GUS with salivary enzymes by co-incubation with 246 bp dsDNA. dsRNA-GUS (2 µg; green arrowhead) incubated in adult *H. halys* saliva with different concentrations of 246 bp dsDNA-AtACT from *A. thaliana* (red arrowhead). After starting the incubation, aliquots were taken at 1, 10, 20, and 30 minutes. 1% SDS was added to stop the reaction (A). dsRNA-GUS incubated in 2 µL of adult *H. halys* saliva. In the ratios, such as 1:2, the first value (1) is indicative of the concentration of dsRNA, while the second value (2) is indicative of the concentration of dsDNA, respectively. The reaction mixtures were incubated at 25 °C and aliquots were taken at increasing times up to 60 minutes and added to 1% SDS to stop the reaction (B). All samples were visualized on a 1% agarose gel with 1xTAE buffer to evaluate the stability of dsRNA-GUS and dsDNA-AtACT (n = 2).

In addition to the 246 bp dsDNA-AtACT from *A. thaliana*, we tested whether the shorter 102 bp dsDNA-*Hh* AGO2 fragment from *H. halys AGO-2* also mediates dsRNA protection from *Hh*NSEs and carried out the degradation experiment under the same experimental conditions (Fig. 6). We tested the stability of 2 µg of dsRNA-GUS when incubated with 2 µL of adult *H. halys* saliva and 2 µg of 102 bp long dsDNA-HhAGO2. Consistent with the results for the 246 bp dsDNA-AtACT, the 102 bp dsDNA-HhAGO2 also reduced the dsRNA-GUS degradation when added to the reaction at equal concentrations (1:1). dsRNA-GUS remained stable up to 60 min after incubation with saliva (Fig. 6C) and showed a distinct band on the gel, but in the identical reaction without dsDNA-HhAGO2, dsRNA-GUS was completely degraded within one minute of treatment with *H. halys* saliva (Fig. 6A).

**Figure 6.**
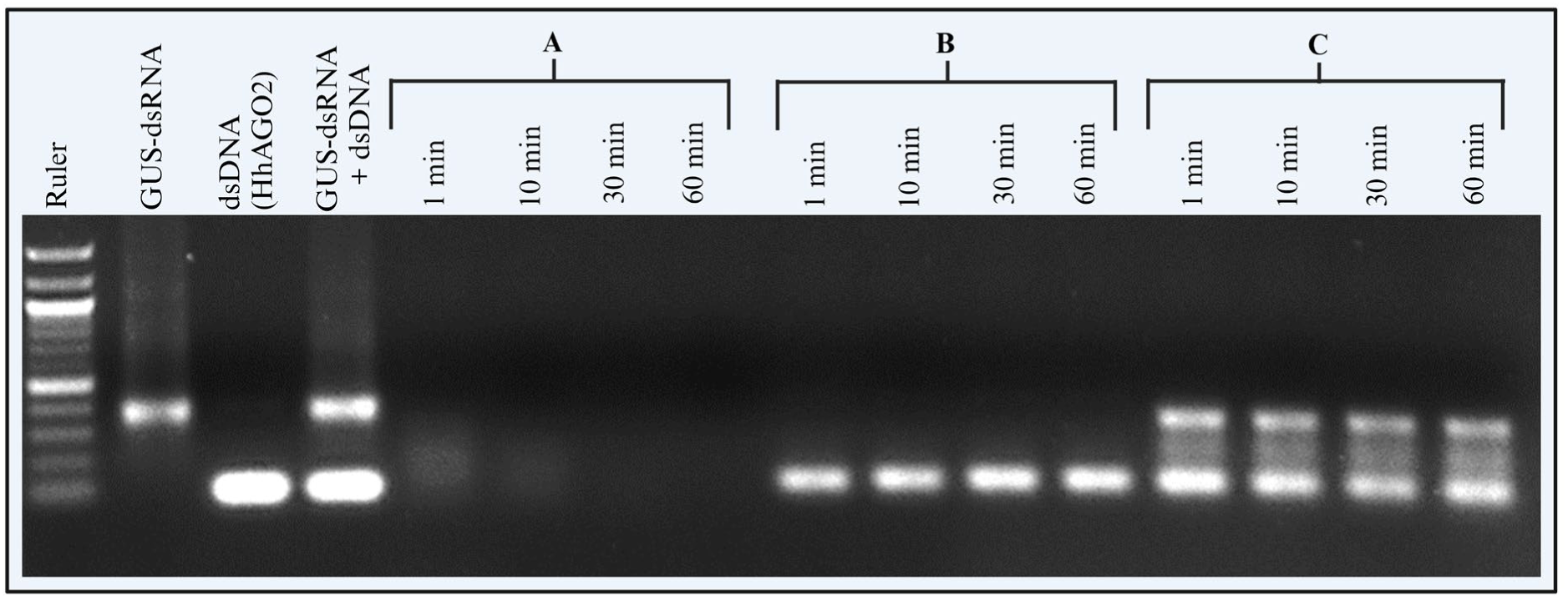
Testing the stability of dsRNA-GUS with saliva by co-incubation with 102 bp dsDNA. dsRNA-GUS (2 µg) was incubated with 2 µL of *H. halys* saliva (A). 102 bp dsDNA-HhAGO2 (2 µg) incubated with 2 µL of *H. halys* saliva (B). dsRNA-GUS (2 µg) with an equal amount of 102 bp dsDNA-HhAGO2 was incubated with 2 µL of *H. halys* saliva (C). After incubation, samples were collected at 1, 10, 30, and 60 min and 1% SDS was added to stop the reaction. The samples were visualized on a 1% agarose gel with 1xTAE buffer to evaluate the stability of dsRNA-GUS and dsDNA-HhAGO2.

Taken together, these results demonstrate the potential of dsDNA as a competitive substrate for dsRNA-degrading nucleases in *H. halys* saliva. We suggest that dsDNA significantly saturates *Hh*NSE, limiting the availability of *Hh*NSEs to degrade dsRNA-GUS. The combined analysis of quantified dsRNA stability on gel images at a 1:1 ratio of dsRNA-GUS to dsDNA-(AtACT/HhAGO2), using the ImageJ tool and SAS 9.0, revealed a significant increase in dsRNA-GUS stability compared to the treatment without dsDNA (F_2,25.1_ = 26.86, p < 0.001) (Fig. S4-6).

**Figure 7.**
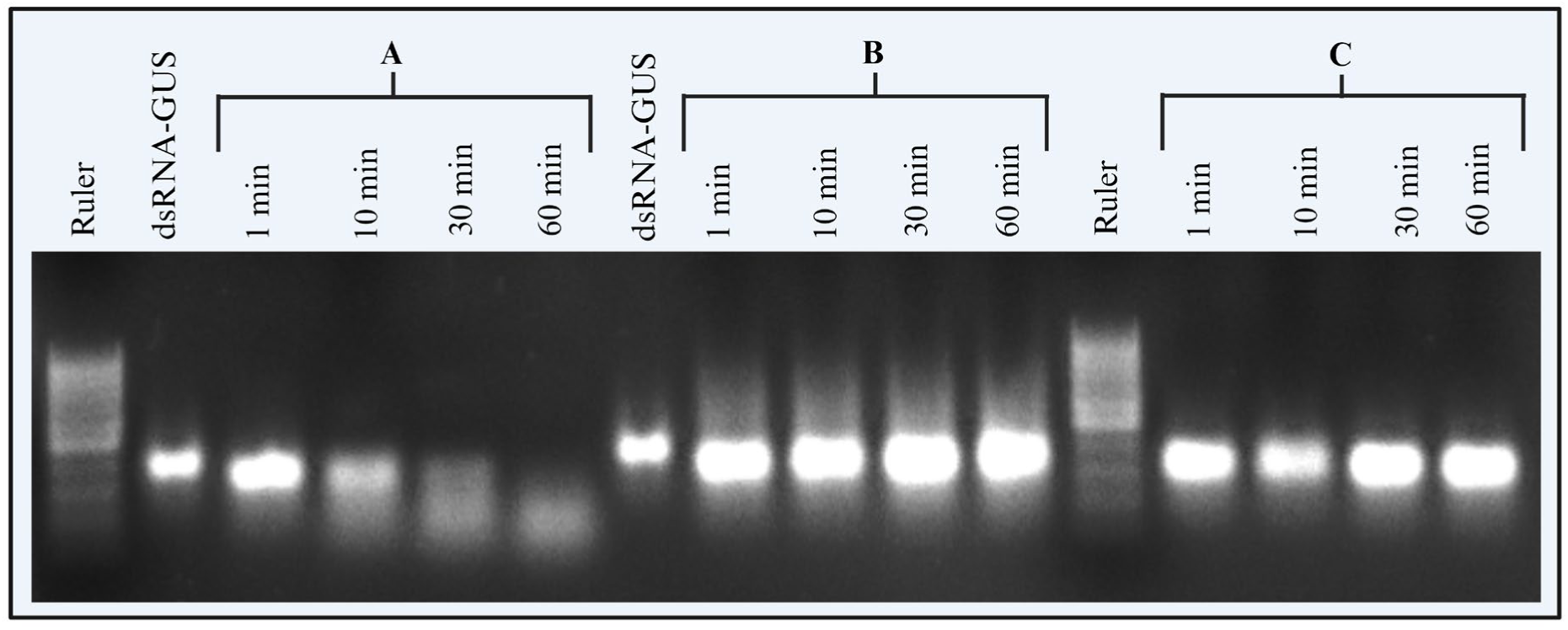
Testing the stability of dsRNA-GUS with saliva from adult *H. halys* by co-incubation with Ultra-Pure Salmon sperm DNA (dsDNA-S). dsRNA-GUS (2 µg) was incubated with 2 µL of *H. halys* saliva (A). dsRNA-GUS (2 µg) with an equal amount of dsDNA-S was incubated with 2 µL of *H. halys* saliva (B). dsRNA-GUS (2 µg) was incubated with 2 µL of Nuclease free water (C). After incubation, samples were collected at 1, 10, 30, and 60 min and 1% SDS was added to stop the reaction. The samples were visualized on a 1% agarose gel with 1xTAE buffer to evaluate the stability of dsRNA-GUS and dsDNA-S (n = 1).

Ultra-pure crude dsDNA-S also demonstrated enhanced stability of dsRNA when incubated with saliva, as illustrated in figure 7. As the results were clearly indicative of enhanced dsRNA stability, this experiment was not replicated. Subsequently, dsDNA-S was used as a co-formulant in dsRNA feeding experiments.

### 3.4 In-vivo dsRNA feeding assays

#### 3.4.1. Use of salmon sperm DNA (dsDNA-S) as a formulation agent to improve oral dsRNA efficacy

To test whether the DNA formulation of dsRNA increases the efficacy of dsRNA-mediated RNAi in live *H. halys*, we examined the effect of orally administered dsRNA-CHC, combined with dsDNA-S, on the silencing of the *HhCHC* gene (Fig. 8). Feeding was confirmed by coloured feeding spots (i.e. salivary sheaths stained with food dye) on parafilm sachets (Fig. S1). Overall, we found differences in gene silencing efficacy between treatments (F_3, 17.1_ = 22.69, p < 0.0001).

**Figure 8.**
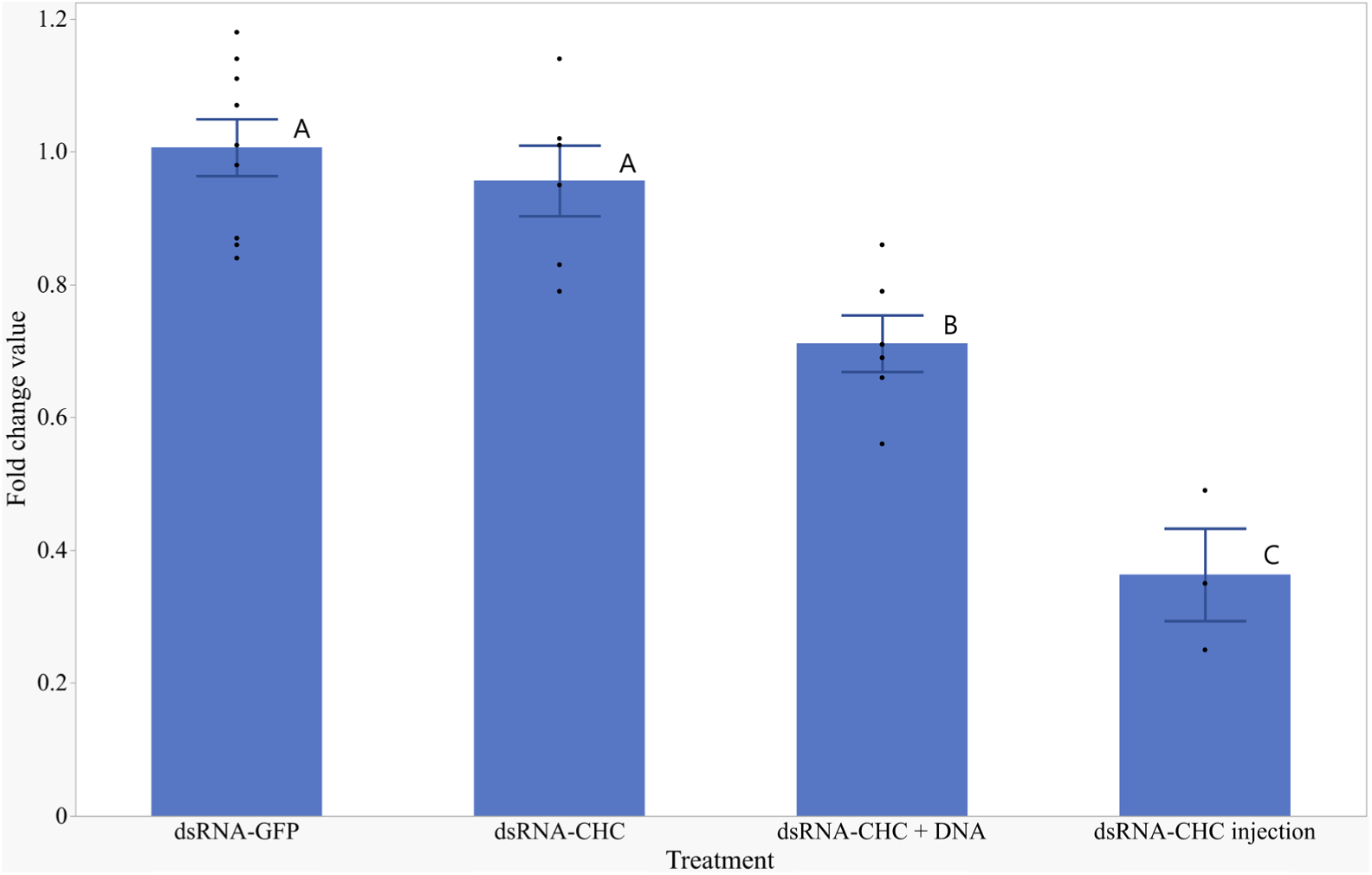
Efficiency of target gene knockdown was evaluated by RT-PCR in dsRNA-fed second instar *H. halys* 72 hours post-treatment. Nymphs were fed 20 µg dsRNA-CHC, 20 µg dsRNA-GFP, or a combination of 20 µg dsRNA-CHC and 20 µg dsDNA-S. As a positive control, 100 ng dsRNA-CHC was injected. Total RNA was converted to cDNA, and mRNA levels were normalized to 18S rRNA. Mean ± SE (n = 3-9) of relative *HhCHC* mRNA levels are shown. Statistical comparisons were made using a Tukey-HSD test.

While *HhCHC* expression levels in insects orally treated with dsRNA-CHC did not differ from insects fed with dsRNA-GFP (p = 0.87), *HhCHC* expression was significantly reduced when dsDNA-S was co-administered with dsRNA-CHC (p = 0.009, dsRNA-CHC + dsDNA-S vs. dsRNA-CHC; p = 0.001, dsRNA-CHC + dsDNA-S vs. dsRNA-GFP). However, direct injection of dsRNA-CHC resulted in an even more pronounced reduction of *HhCHC* expression, approximately 6-fold compared to oral dsRNA-GFP (p < 0.0001) and dsRNA-CHC (p < 0.001), and 3-fold compared to oral administration of dsRNA-CHC combined with dsDNA-S (p = 0.02), all post-hoc comparisons Tukey-HSD.

### 3.5 dsRNA-CHC injection significantly increases mortality in *H. halys* compared to controls

To assess the impact of dsRNA-CHC on the survival of *H. halys*, we selected second-stage insects and injected 100 nL (1 µg/µL concentration) of dsRNA per insect. Survival data were recorded over a 13-day period following the injection.

We observed high mortality upon the injection of dsRNA-CHC close to 100 percent (least square mean 93.75% ± 6.83 SE, Fig. 9). In all control treatments (dsRNA-GFP, NF water, untreated), mortality only ranged around 25% (Least square means dsRNA-GFP: 19.39 ± 6.83, NF water: 19.81 ± 6.83, untreated: 28.81 ± 6.83). Statistical analysis of our data clearly revealed differences across the treatments (GLM, F_3, 189_ = 12.43, p < 0.001). Specifically, the injection of dsRNA-CHC caused substantially higher mortality compared to the other treatments (p < 0.001 for all comparisons), that were not different from each other (p > 0.823).

**Figure 9.**
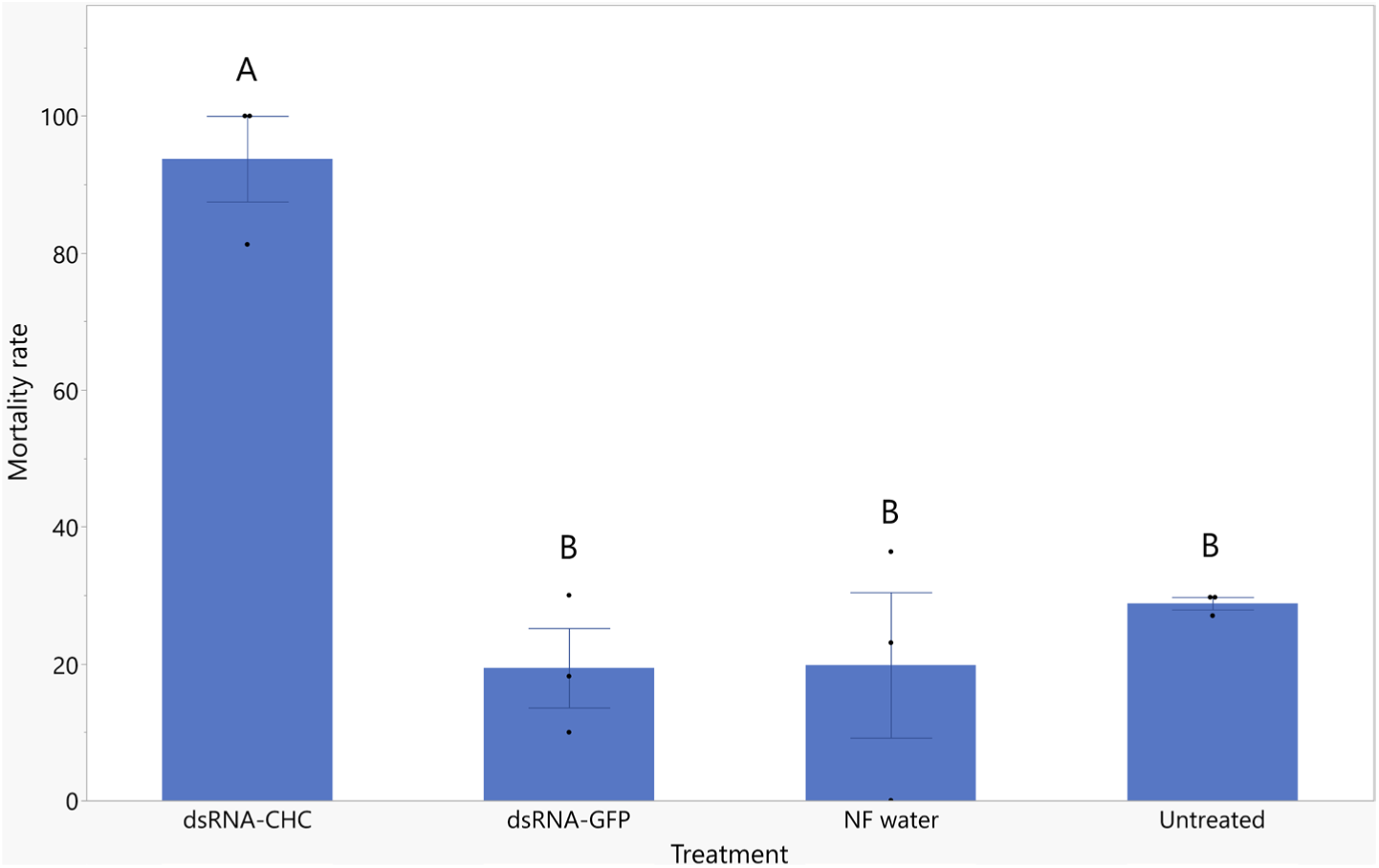
Mortality rates of second instar *H. halys* nymphs injected with dsRNA-CHC, dsRNA-GFP, NF water, or left untreated groups. In total, three experiments were performed, with sample sizes ranging from 10 to 37 individuals per treatment. Mortality was assessed over 13 days and analysed by ANCOVA using the standard least squares method, followed by a Tukey-HSD post-hoc test. Mean ± SE (n = 10-37) of cumulative mortality across experiments is shown. Treatments with different letters indicate significant differences (Tukey-HSD, p < 0.001).

### 3.6 Survival of *H. halys* following oral ingestion of dsRNA-CHC with and without dsDNA

To evaluate the effect of adding dsDNA to dsRNA-CHC on *H. halys* survival via oral delivery, we fed second-stage insects 200 µL of dsRNA (100 ng/µL) to groups of five. Survival was monitored over 14 days while they fed on fresh green bush beans post-treatment.

No significant differences were observed across all treatments (GLM, χ²(6) = 6.6784, p = 0.3516), indicating that the oral delivery of dsRNA-CHC did not result in increased mortality, irrespective of the addition of dsDNA (Fig. S3).

## 4. Discussion

RNA interference (RNAi) has emerged as a promising strategy for pest control, offering specificity and reduced environmental impact compared to traditional chemical pesticides. Our study addresses one of the major challenges in the application of RNAi in agriculture - the degradation of dsRNA by nucleases exists in the target pests. We found that both saliva and salivary gland extracts of *H. halys* rapidly degrade dsRNA *in vitro*, a finding that is consistent across all tested developmental stages of the pest. Furthermore, our data suggest that HhNSE is a key enzyme involved in this degradation process. Our main finding is that dsDNA acts as a competitive inhibitor to protect dsRNA from enzymatic degradation, significantly increasing its stability and the efficacy of gene silencing.

dsRNases impair the oral delivery efficiency of dsRNA in many insects, including *H. halys*, thereby reducing the effectiveness of RNAi-based pest control strategies^9,48^. Notably, previous research documented significant dsRNA degradation in the saliva of adult *H. halys*^27^. However, it remained uncertain whether this degradation extends to other developmental stages. Our demonstration of dsRNase activity in saliva, salivary gland extracts and hemolymph from different developmental stages of *H. halys* may support future strategies for effective control of this important insect pest.

Our ex vivo dsRNA stability assays in *H. halys* have shown that dsRNA exhibits prolonged stability in the hemolymph, exceeding 4 hours, in contrast to its rapid degradation in saliva at all growth stages tested (Fig. 1 & 7). This observation is consistent with the other insects showing RNAi susceptibility patterns when dsRNA injected directly into the hemolymph^44,49^. In such insects, dsRNA remains intact and functional when injected into the hemolymph, thereby ensuring dsRNA to remain active long enough to be taken up by cells and activate the RNAi mechanism, resulting in target gene silencing. The constancy of this pattern across different RNAi-sensitive insect species highlights the significance of dsRNA stability in the efficacy of RNAi.

Conversely, the rapid destruction of dsRNA observed in *H. halys* saliva is consistent with findings in RNAi-resistant insects^15,17,31,50,51^. Salivary nucleases rapidly degrade dsRNA in these species, preventing the dsRNA from entering into the cells and silencing target genes, leading to ineffective oral-route RNAi. For example, in *Nezara viridula* where severe nuclease activity observed in saliva, silencing NSE improved oral RNAi efficacy to 65% mortality compared to 43.3% without NSE silencing when treated with dsRNA targeting the *alpha coatomer* gene^34^. We found that *HhNSE* has significantly higher expression levels in salivary glands compared to other known insect dsRNase’s such as *HhEri-1* and *HhSDN-1*^17,52,53^, suggesting a predominant role of *HhNSE* in salivary dsRNA digestion. This is supported by literature data demonstrating that silencing of *NSE* in other species such as *Spodoptera frugiperda*^54^, *N. viridula*^34^, *Bemisia tabaci*^55^, and *Tribolium castaneum* ^56^ enhances the efficacy of RNAi.

Given the observed broad substrate specificity of NSEs^29,30^, we hypothesised that dsDNA could act as a competitive inhibitor for salivary HhNSE and thereby mitigate dsRNA degradation in *H. halys*. Our results (Fig. 6& 7) clearly demonstrate that the addition of dsDNA to saliva samples significantly reduces dsRNA degradation, marking the first instance of using DNA as a competitive inhibitor against dsRNA-degrading nucleases. It would be beneficial to further characterise the principle of competitive inhibition by producing and purifying the enzyme in order to test the efficiency of dsDNA in protecting dsRNA from HhNSE-mediated degradation in the future.

The in vivo results demonstrate that the presence of dsDNA-S not only enhances the stability of dsRNA in ex vivo conditions but also amplifies RNAi efficacy in vivo (Fig. 8). This is indicated by the increased efficacy of target gene silencing achieved by ingesting both dsDNA-S and dsRNA-CHC together. Nevertheless, the efficacy of *HhCHC* silencing achieved by the oral delivery of dsDNA-S in conjunction with dsRNA-CHC is considerably lower to that observed following dsRNA-CHC injection. The direct injection of dsRNA-CHC into the hemocoel resulted in complete mortality within 13 days, a finding that is consistent with previous studies on Pentatomidae insects, such as *N. viridula* ^34^. In contrast, the oral delivery of dsRNA-CHC combined with dsDNA-S resulted in lower gene silencing levels that did not induce mortality in H. halys. Nevertheless, the RT-PCR results indicating a considerable enhancement in target gene silencing imply that the incorporation of dsDNA as a co-formulant to competitively inhibit NSE represents a promising strategy. Nevertheless, it is essential to address the other possible biological barriers that impede the delivery of dsRNA in order to optimise this approach.

The potential biological barriers to dsRNA delivery in *H. halys* can be broadly categorised as follows: (a) dsRNase activity in the oral tract, (b) dsRNA uptake and release in epithelial cells, (c) functionality of the RNAi machinery, and (d) intercellular transport of dsRNA. Our injection assays demonstrated that barriers c and d are not present, while barrier a can be overcome by our DNA formulation. Therefore, the remaining barrier b may have restricted the efficient oral delivery of dsRNA. A more profound comprehension of the mechanisms governing dsRNA uptake and release in gut epithelial cells is vital for the development of more efficacious dsRNA delivery systems in *H. halys*.

Reason for the lower silencing observed with dsDNA plus dsRNA feeding in *H. halys* can be related to difficulties with dsRNA uptake into midgut cells. Beyond the endocytic dsRNA uptake mechanism in midgut cells, SID-like proteins have been shown to play a crucial role in dsRNA uptake in several insects^57^. In the hemipteran insect *Nilaparvata lugens*, a SID-like protein was shown to be involved in dsRNA uptake and to increase the efficiency of systemic RNAi ^58^. However, as Sparks M et al. 2014^59^ have indicated, SID or SID-like protein is not present in the *H. halys* transcriptome, which provides further evidence of the potential difficulties associated with dsRNA uptake. Nevertheless, it would be appropriate to test whether there are indeed difficulties in dsRNA uptake in midgut cells before reaching a conclusion that the lack of SID protein hinders RNAi efficiency.

Another interesting finding was that the dsDNA was not completely degraded by the salivary enzymes (Fig. 5 & 6), similar to the observation in *Lygus lineolaris*^51^ where separate incubation of a similar amount of dsDNA and dsRNA with saliva did not completely degrade the dsDNA but only the dsRNA. It is possible that HhNSE has a high affinity for degrading dsRNA compared to dsDNA, a pattern reported in *Apolygus lucorum*^60^. Also, in the study by Meiss et al., 1999^29^ documented that NSEs in *B. mori* showed a predilection for degrading dsRNA over other nucleic acid forms in the insect body. An alternative explanation for the observed results is that the dsDNA may have saturated the available HhNSE, allowing the remaining dsRNA to remain intact and to be protected from HhNSE for some time. This would allow the midgut epithelial cells to complete its uptake of the free dsRNA.

To ensure the durability of dsRNA after oral administration, several studies have successfully used metal ion chelators such as ethylenediaminetetraacetic acid (EDTA)^14^, chitosan-based nanoparticles^61,62^, and lipid molecules such as Lipofectamine 2000^14^ to protect dsRNA from degradation by insect nucleases or to increase RNAi efficiency^63^. However, using conventional nuclease inhibitors such as EDTA, SDS could also interfere with the efficiency of key RNAi-associated nucleases such as Dicer enzymes (acting as dsRNA endoribonucleases) and Argonaute proteins (RNA-guided endonuclease) upon entering into the epithelial cells of insect gut. Moreover, cytotoxicity to non-target organisms and later market approval of nanoparticle-based formulation needs to be concerned^64–66^. In contrast, dsDNA has been identified as a biologically safe co-formulant, capable of enhancing the efficiency of RNAi pesticides without, or with minimal, negative environmental impact. Presence of dsDNA in the diets of practically all living organisms, it is anticipated that feeding dsDNA may pose no or low risks depending on the doses administered^67^. Nevertheless, it is important to test the environmental and ecological safety of introducing dsDNA into ecosystems even though it is already widespread in natural environment. Further research is essential to optimise and develop comprehensive formulations incorporating DNA.

Limitations associated with the use of dsDNA as a co-formulant could encompass the potential activation of the immune system when dsDNA is administered at high concentrations^68–70^. Additionally, and of significant concern, is the possibility that dsDNA may compete with dsRNA for uptake and mobility within gut tissues, potentially compromising the efficacy of dsRNA delivery. Therefore, it is crucial to optimize the concentrations of both nucleic acids to fully harness the advantages of this method in enhancing oral RNAi efficiency in *H. halys* and other insect pests representing similar challenges.

Historical applications of DNA, including the use of salmon sperm DNA in environmental RNA extractions and as a blocking agent in hybridization techniques^71,72^, have demonstrated the utility of DNA in nucleic acid isolation and hybridization technologies. Our use of DNA to enhance dsRNA stability suggests a potential for improved RNAi-based pest control strategies.

## 4. Conclusion

In conclusion, our ex vivo analysis revealed significant nuclease activity in the saliva of *H. halys* across various developmental stages tested. Furthermore, RT-PCR data confirmed high expression of *HhNSEs* in the salivary glands. The addition of dsDNA as a formulant proved effective in reducing dsRNA degradation by these salivary enzymes when incubated ex vivo. These findings were further supported by in vivo assays, which demonstrated that the co-delivery of dsRNA with dsDNA significantly enhanced gene silencing compared to dsRNA alone. Although no significant increase in mortality rates was observed with the incorporation of the dsDNA-S formulation into the dsRNA-CHC, it is hypothesised that there may be a lack of an efficient dsRNA uptake mechanism in *H. halys*. This therefore suggests the need for further research to gain a deeper understanding of the dsRNA uptake mechanism in *H. halys* into gut epithelial cells. These results offer valuable insights into optimizing dsRNA delivery in *H. halys* and highlight potential broader applications for overcoming non-specific nuclease-related difficulties to dsRNA delivery in other contexts.

## Supporting information

ex_vivo_data

## Acknowledgements

The authors express gratitude to Robert Bischoff (Department of Applied Entomology, University of Hohenheim) for his invaluable assistance with the modelling and statistical analysis of quantified agarose gel data using SAS software. Special thanks to Anja Kuehne for her contribution to insect rearing and to Swathi Ramapuram for providing technical support during the *ex-vivo* dsRNA stability assays. We would like to express our sincerest gratitude to the funding agencies for this project, without whom it would not have been possible. This project was funded by QS Qualität und Sicherheit GmbH (QS), the Ministry of Land Use, Development, and Environment (MLR) of Germany, and the Federal Ministry for Economic Affairs and Energy (BLE).

## Conflict of interest

The authors declare that the research was conducted in the absence of any commercial or financial relationships that could be construed as a potential conflict of interest.

## Author contribution

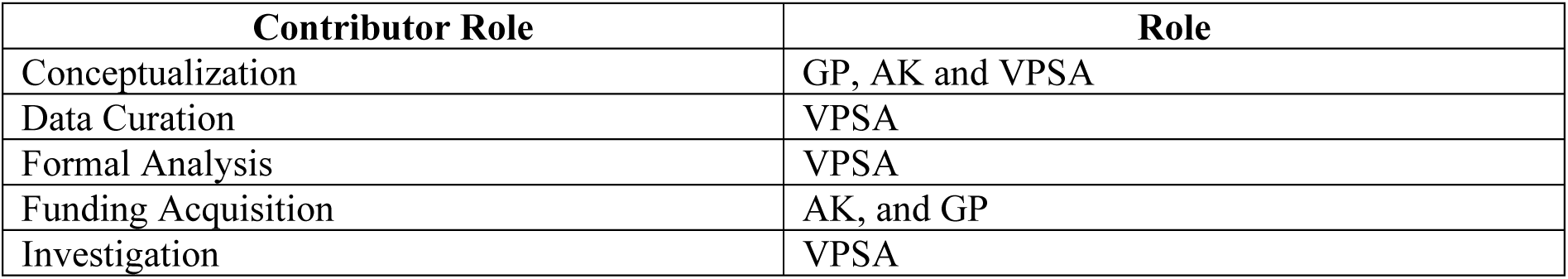
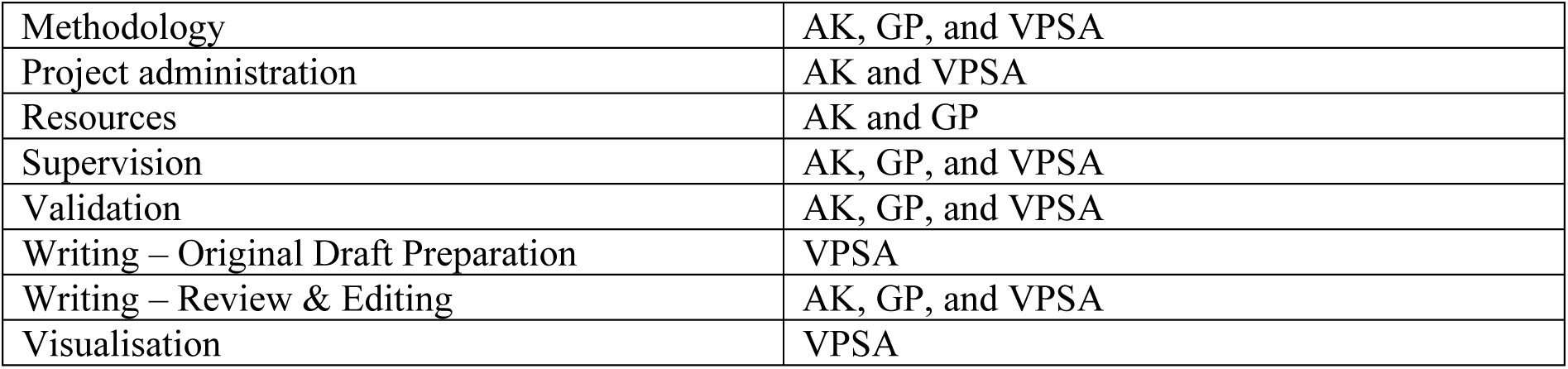

## Data availability

All datasets generated for this study are included in the Supplementary Material.

## Notes

### Competing Interest Statement

The authors have declared no competing interest.

