## Supplementary Material for "Double-stranded DNA reduces dsRNA degradation in the saliva and significantly enhanced RNAi-mediated gene silencing in *Halyomorpha halys*"

---

#### 1 Primers and dsRNA sequences used in this study:

| Gene name | Accession id | Primer sequence (5' – 3') | Product size (bp) |
|---|---|---|---|
| Actin-2 (*Arabidopsis thaliana*) | NM_001338359.1 | Fw: GGAAGGATCTGTACGGTAAC<br>Rv: TGTGAACGATTCCTGGACCT | 246 |
| Argonaute-2 | XM_024358504.1 | Fw: AGTTGGTTCAGTGGAGAGTTG<br>Rv: TGCCACATTCCCTTCCATAAT | 102 |
| Clathrin heavy chain | XM_014431604.1 | Fw: CTGATTGGCTTGTGGGATACT<br>Rv: GTCAGCTGTTCGTGGTACTT | 139 |
| Exo-ribonuclease-1 | XM_024361541.1 | Fw: GCAGAGGAGTCGAAGGTTTAG<br>Rv: GTCAGTGTCAGGTAGGGTTTC | 91 |
| Small RNA degrading nuclease 1 (Exo nuclease) | XM_014423854.2<br>XM_014423853.2 | Fw: GGCTGTTGACTGTGAGATGT<br>Rv: GGATTGTAGGGCTTCACTAAGG | 114 |
| DNA/RNA non-specific endonuclease isoform 2 | XM_024362815.1 | Fw: CGTCACCGGAACAGAAGGAA<br>Rv: AATGCAGGAGTCGTCTTGGG | 136 |

**Supplementary table 1.** primers used in this study.

| | dsRNA Sequence |
|---|---|
| dsRNA-GUS | CTCTACACCACGCCGAACACCTGGGTGGACGATATCACCGTGGTGACGCATGTCGCGCAAGACTGTAACCACGCGTC<br>TGTTGACTGGCAGGTGGTGGCCAATGGTGATGTCAGCGTTGAACTGCGTGATGCGGATCAACAGGTGGTTGCAACTG<br>GACAAGGCACTAGCGGGACTTTGCAAGTGGTGAATCCGCACCTCTGGCAACCGGGTGAAGGTTATCTCTAT |
| dsRNA-GFP | TGATCGCGCTTCTCGTTGGGGTCTTTGCTCAGGGCGGACTGGGTGCTCAGGTAGTGGTTGTCGGGCAGCAGCACGGG<br>GCCGTCGCCGATGGGGGTGTTCTGCTGGTAGTGGTCGGCGAGCTGCACGCTGCCGTCCTCGATGTTGTGGCGGATCTT<br>GAAGTTCACCTTGATGCCGTTCTTCTGCTTGTCGGCCATGATATAGACGTTGTGGCTGTTGTAGTTGTACTCCAGCTTG<br>TGCCCCAGGATGTTGCCGTCCTCCTTGAAGTCGATGCCCTTCAGCTCGATGCGGTTCACCAGGGTGTCGCCCTCGAAC<br>TTCACCTCGGCGCGGGTCTTGTAGTTGCCGTCGTCCTTGAAGAAGATGGTGCGCTCCTGGACGTAGCCTTCGGGCATG<br>GCGGACTTGAAGAAGTCGTGCTGCTTCATGTGGTCGGGGTAGCGGCTGAAGCACTGCACGCCGTA |
| dsRNA-CHC | TTGTTTTGCTTTGCTGTTCGTACACTTCAAGGAGGAAAGCTTCATATAATTGAGGTTGGACAGCCTCCTACAGGAAAC<br>CAACCATTCTCAAAAAAAGCAGTGGATGTATTTTTTCCAGTAGAAGCCCAAAATGATTTTCCAGTTGCAATGCAGGT<br>TAGCTCTAAATATGATGTAATCTATCTTATTACAAAGTATGGATACATTCATTTGTATGATTTGGAAACAGCTACATG<br>TATTTACATGAACCGTATTAGTATTGATACTATATTTGTAACCGCTCCTCATGAATCAACTGGTGGTATCATAGGTGT<br>GAATAGAAAAGGCCAGGTGTTATCAGTGAGTGTTGAAGAGGACCATATAATCCCATATATCAATAATATATTACAAA<br>ATCCTGATCTAGCATTACGCATGGCTG |

#### 2 Supplementary figures

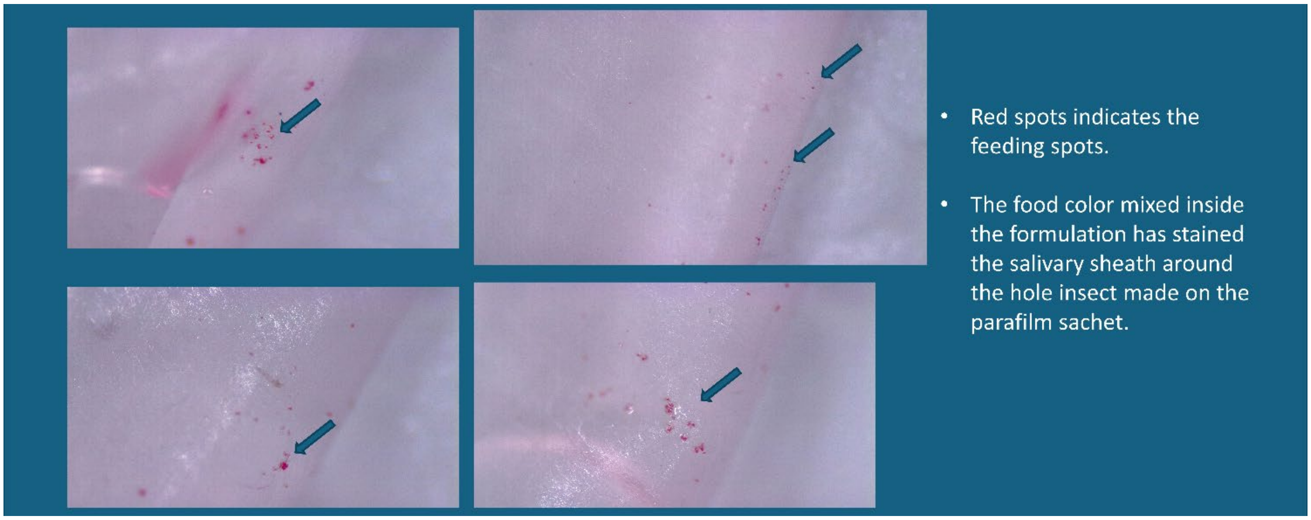

**Supplementary figure 1. Feeding sites of *Halyomorpha halys* on parafilm sachet are stained with red food color (Liebensmittel Farbe - Rot, Dr. August Oetker Nahrungsmittel KG, Germany). The arrows point to the feeding sites where the protein-rich salivary sheath was stained with the red food color, which was previously mixed with the sugar solution packed inside.**

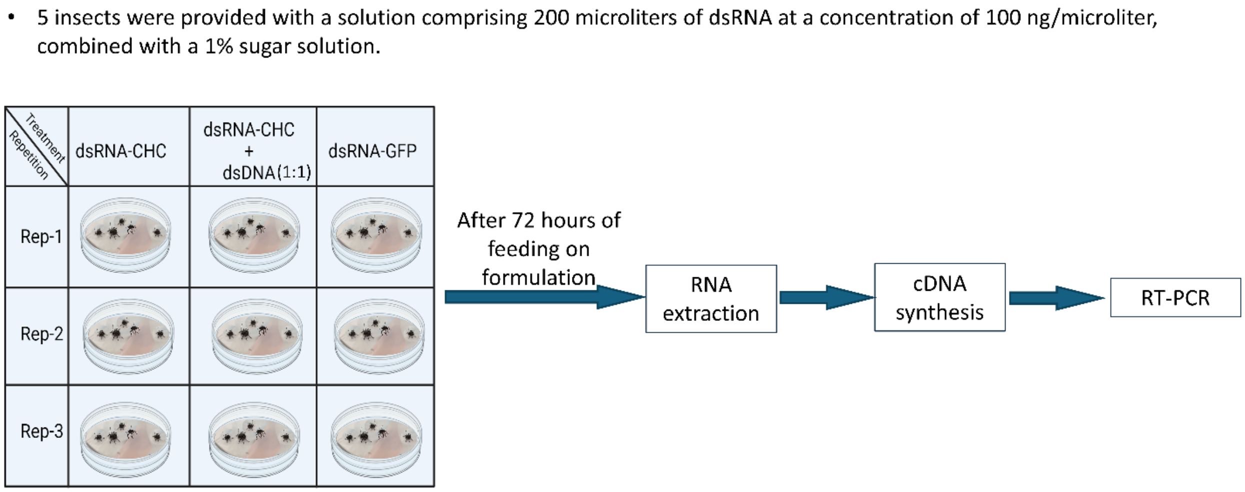

**Supplementary figure 2. Experimental design of dsRNA feeding assays with *H. halys* nymphs. A 200 microliters of a dsRNA solution with a concentration of 100 ng/microliter were given to five insects along with a 1% sugar solution. Each sample includes five insects. Three combinations of dsRNA-CHC, dsRNA-CHC with dsDNA at a 1:1 ratio, and dsRNA-GFP were examined. Rep-1, Rep-2, and Rep-3 were the three replications of each treatment. Following a 72-hour feeding**

**period, the insects' RNA was extracted, and then cDNA was synthesized and subjected to RT-PCR analysis.**

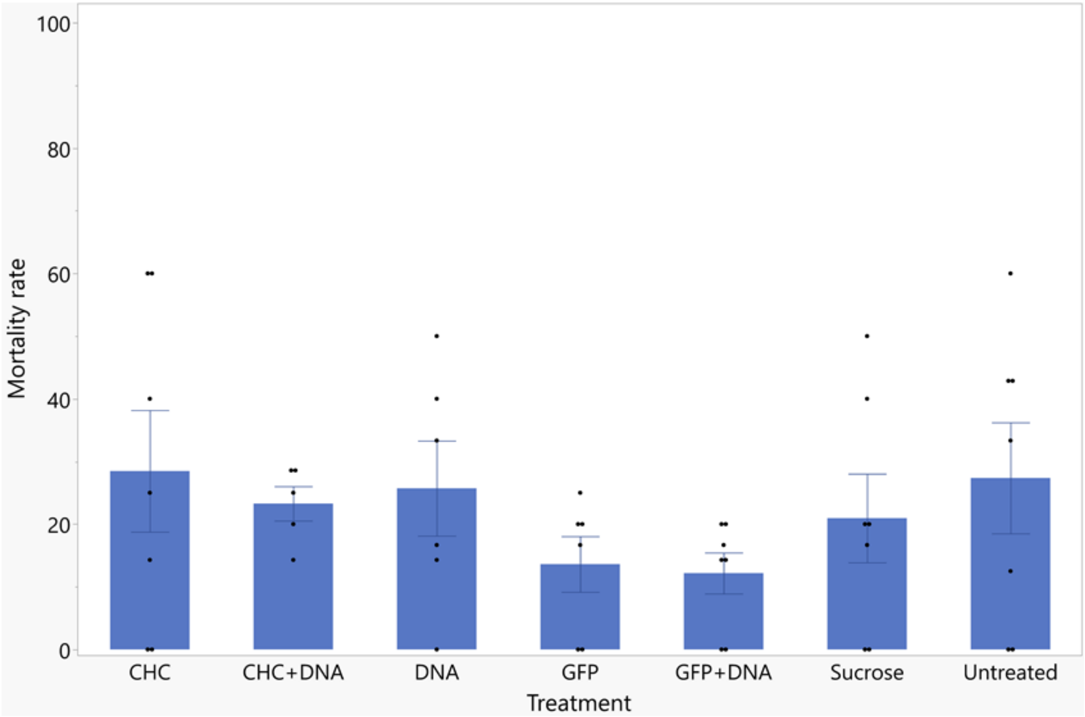

**Supplementary figure 3. Mortality of second instar *H. halys* nymphs was assessed over a 14-day period following oral delivery of the dsRNA treatments. Nymphs were fed 200 µL of 100 ng/µL dsRNA-CHC, dsRNA-CHC plus dsDNA-S, dsRNA-GFP, dsRNA-GFP plus dsDNA-S, dsDNA-S, 1% sucrose, and untreated control. A 1% sucrose solution and an untreated control group were also included. Survival was recorded daily after 72 hours of feeding on dsRNA solutions. Fresh green beans were provided after dsRNA feeding. Mean ± SE (n = 5-7) are shown. Error bars indicate the standard error of the mean (SEM). Dots represent individual replicates, and outliers are visible for certain treatments. Statistical analysis was performed using the GLM.**

#### 3 dsRNase's expression in salivary glands - RT-PCR assay:

Analysis of RT-PCR data

Gene expression ratio = $(E_{GOI})^{\Delta Ct\ GOI} / (E_{GOI})^{\Delta Ct\ CG}$

E= (Primer efficiency/100) +1

GOI = Gene of Interest

CG = calibrator gene

| **Primer efficiencies** | **Efficiency %** | **Converted efficiency E** | |
|---|---|---|---|
| 18s rRNA | 89.8 | 1.898 | |
| 60 SRP | 93.7 | 1.937 | |
| dsRNase | 92.5 | 1.925 | |
| Eri 1 | 100.7 | 2.007 | |
| SDN | 97.1 | 1.971 | |
| **Target gene** | **Avg Ct** | **ΔCt (ct target - control average)** | **Gene expression ratio** |
| dsRNase | 20.05356689 | 5.765001728 | 108.9566518 |
| dsRNase | 17.08789167 | 8.730676948 | 180.721711 |
| dsRNase | 17.11952286 | 8.699045762 | 200.9224265 |
| Eri 1 | **27.13244576** | -1.313877142 | 1 |
| Eri 1 | **25.07071046** | 0.747858161 | 1 |
| Eri 1 | **25.25254964** | 0.566018981 | 1 |
| SDN | 24.45775319 | 1.36081543 | 6.28816867 |
| SDN | 23.44744578 | 2.371122838 | 2.968080417 |
| SDN | 23.43285525 | 2.385713369 | 3.402438341 |

**Supplementary table 2 The data presented here are the raw data used for the analysis of Cq values from RT-PCR experiments. The table includes the primer efficiencies employed for data analysis using the Pfaffl method. It presents the efficiency percentages of the primers and their corresponding converted efficiencies (E), as well as the average Cq values, ΔCq (Cq target - control average), and gene expression ratios for each target gene.**

### 4 Raw images of agarose gels used produced in this study

#### 4.1.1 Saliva

##### 4.1.1.1 dsRNA degradation assay with saliva- dsDNA-AtACT (different DNA concentrations)

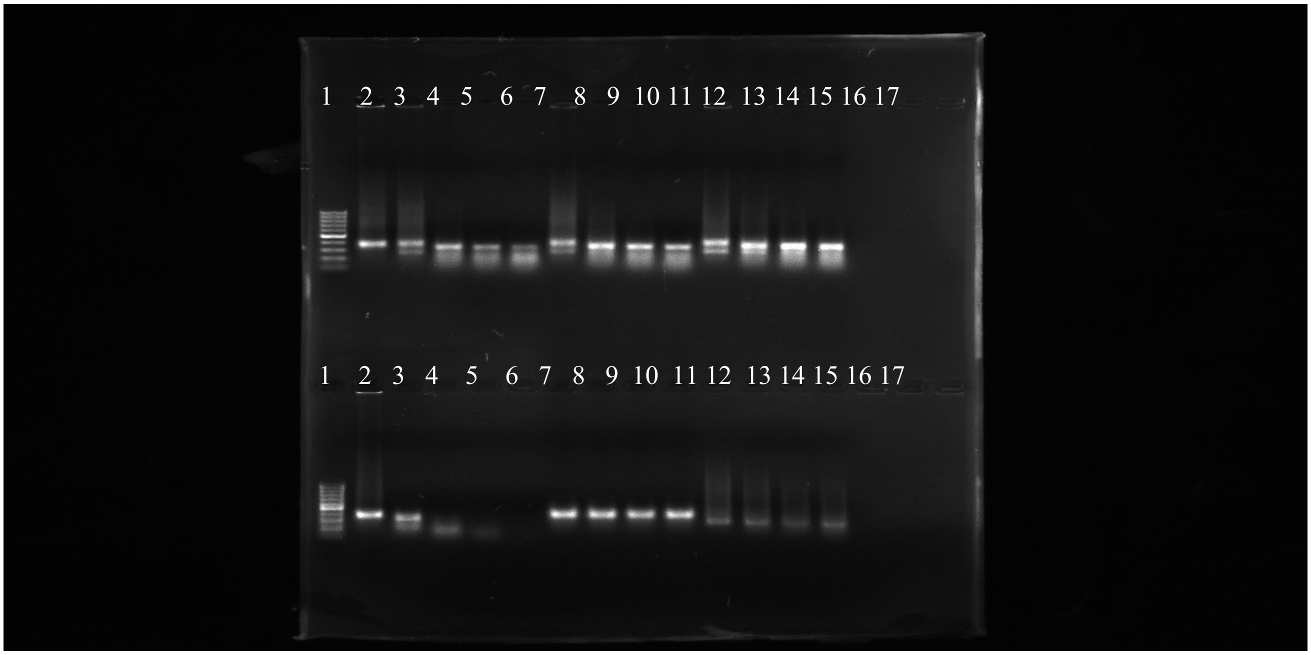

| well | sample name | Treatment | Well | Sample name | Treatment |
|---|---|---|---|---|---|
| 1 | Ladder | 4 µg dsGUS + 2 µg dsDNA + 2 µL saliva(adults) | 1 | Ladder | |
| 2 | dsRNA-GUS | | 2 | dsRNA-GUS | 2 µg dsGUS + 2 µL saliva |
| 3 | 1 min | | 3 | 1 min | |
| 4 | 10 min | | 4 | 10 min | |
| 5 | 30 min | | 5 | 30 min | |
| 6 | 60 min | | 6 | 60 min | |
| 7 | 1 min | 2 µg dsGUS + 3 µg dsDNA + 2 µL saliva(adults) | 7 | 1 min | 2 µg dsGUS + 2 µL water |
| 8 | 10 min | | 8 | 10 min | |
| 9 | 30 min | | 9 | 30 min | |
| 10 | 60 min | | 10 | 60 min | |
| 11 | 1 min | 2 µg dsGUS + 4 µg dsDNA + 2 µL saliva(adults) | 11 | 1 min | 4 µg dsDNA + 2 µL saliva |
| 12 | 10 min | | 12 | 10 min | |
| 13 | 30 min | | 13 | 30 min | |
| 14 | 60 min | | 14 | 60 min | |
| 15 | | | 15 | | |
| 16 | | | 16 | | |
| 17 | | | 17 | | |

#### 4.1.1.2 Saliva - 1:0.5 dsDNA-AtACT:dsRNA degradation assay -

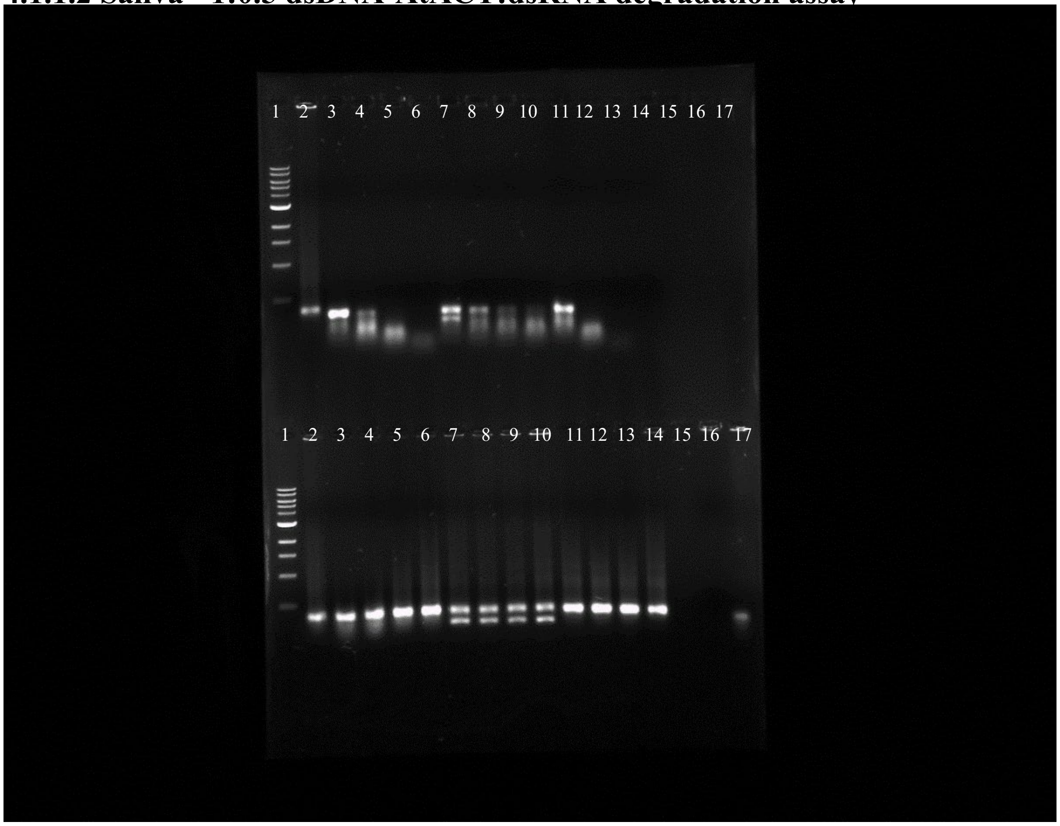

| well | sample name | Treatment | Well | Sample name | Treatment |
|---|---|---|---|---|---|
| 1 | Ladder | | 1 | Ladder | |
| 2 | dsRNA-GUS | | 2 | dsRNA-GUS | |
| 3 | 1 min | 4 µg dsRNA-GUS + 2 µL saliva (adults) | 3 | 1 min | 4 µg dsRNA-GUS + 2 µL water |
| 4 | 10 min | | 4 | 10 min | |
| 5 | 30 min | | 5 | 30 min | |
| 6 | 60 min | | 6 | 60 min | |
| 7 | 1 min | 2 µg dsRNA-GUS + 1 µg dsDNA-AtACT + 2 µL saliva (adults) | 7 | 1 min | 2 µg dsRNA-GUS + 1 µg dsDNA-AtACT + 2 µL water |
| 8 | 10 min | | 8 | 10 min | |
| 9 | 30 min | | 9 | 30 min | |
| 10 | 60 min | | 10 | 60 min | |
| 11 | 1 min | 2 µg dsRNA-GUS + 1 µL RiboLock + 2 µL saliva (adults) | 11 | 1 min | 2 µg dsRNA-GUS + 1 µL RiboLock + 2 µL water |
| 12 | 10 min | | 12 | 10 min | |
| 13 | 30 min | | 13 | 30 min | |
| 14 | 60 min | | 14 | 60 min | |
| 15 | | | 15 | | |
| 16 | | | 16 | | |
| 17 | | | 17 | | |

##### 1.1.1.3. **dsRNA degradation assay - saliva**

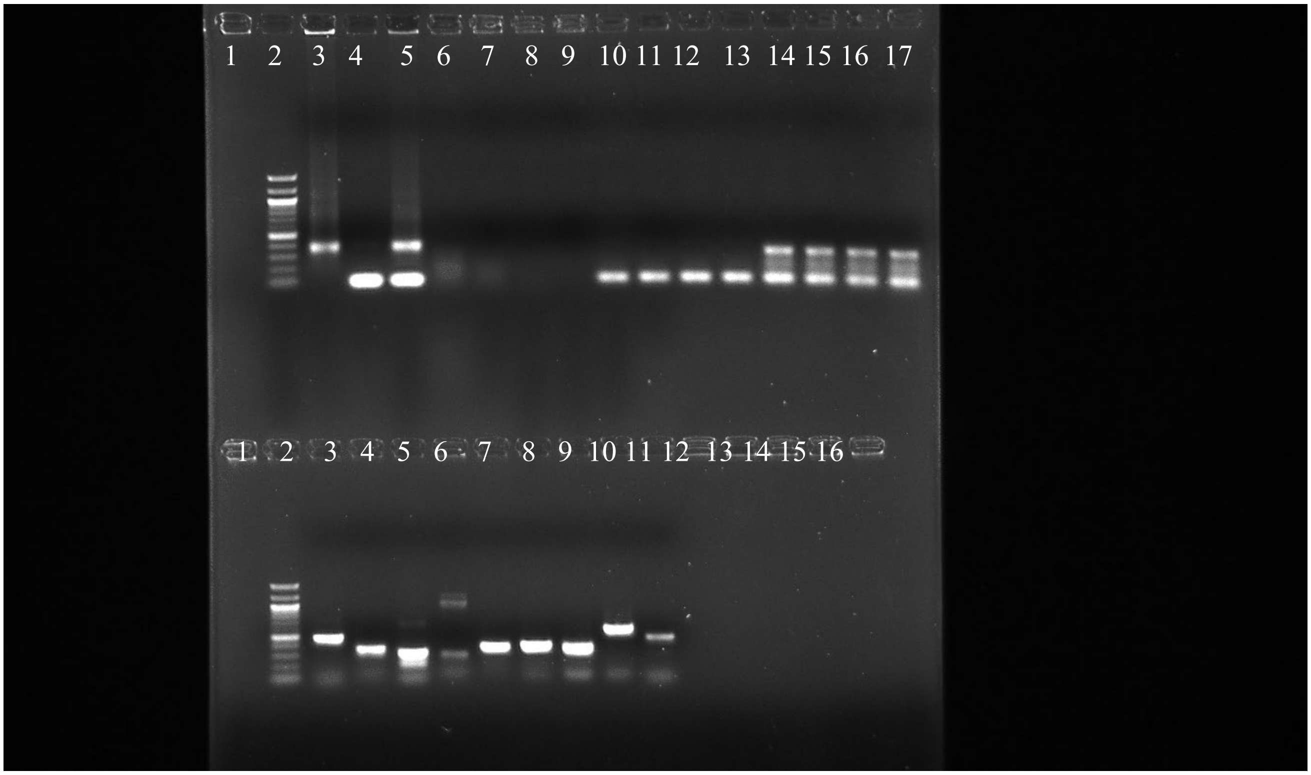

| well | sample name | Treatment | Well | Sample name | Treatment |
|---|---|---|---|---|---|
| 1 | | | 1 | | |
| 2 | Ladder | | 2 | | |
| 3 | dsRNA-GUS | 4 µg dsGUS + 2 µL saliva(adults) | 3 | | |
| 4 | dsDNA-HhAGO2 | | 4 | | |
| 5 | dsRNA-GUS+dsDNA-HhAGO2 | | 5 | | |
| 6 | 1 min | | 6 | | |
| 7 | 10 min | | 7 | | |
| 8 | 30 min | | 8 | | |
| 9 | 60 min | | 9 | | |
| 10 | 1 min | 2 µg dsDNA + 2 µL saliva(adults) | 10 | | |
| 11 | 10 min | | 11 | | |
| 12 | 30 min | | 12 | | |
| 13 | 60 min | | 13 | | |
| 14 | 1 min | 2 µg dsGUS + 2 µg dsDNA + 2 µL saliva(adults) | 14 | | |
| 15 | 10 min | | 15 | | |
| 16 | 30 min | | 16 | | |
| 17 | 60 min | | 17 | | |

#### 1.1.1.4. ***H.halys*** **DNA and dsRNA degradation assay - saliva**

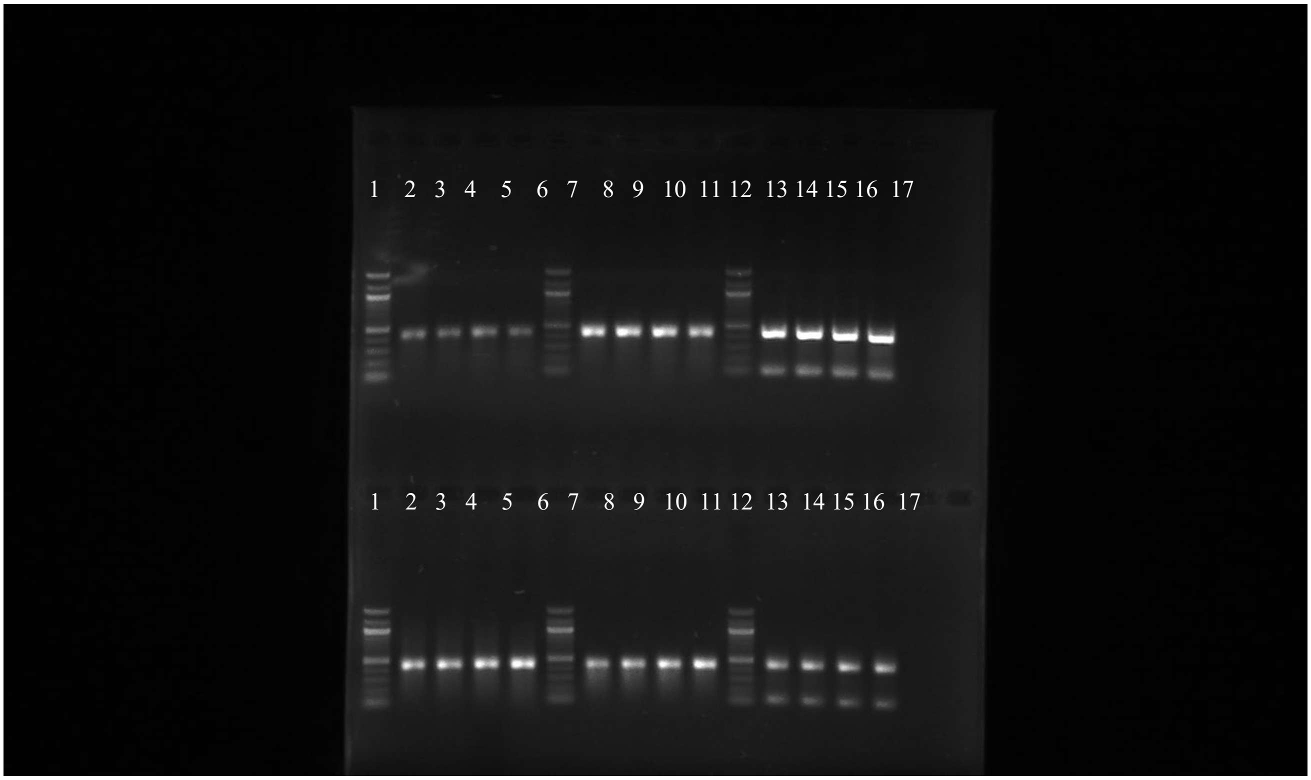

| well | sample name | Treatment | Well | Sample name | Treatment |
|---|---|---|---|---|---|
| 1 | Ladder | | 1 | Ladder | |
| 2 | 1 min | 2 µg dsRNA-Ache + 2 µL saliva (adults) | 2 | 1 min | 4 µg dsRNA-Ache + 2 µg salmon dsDNA + 2 µL saliva (adults)<br><br>-Not considered for analysis- |
| 3 | 10 min | | 3 | 10 min | |
| 4 | 30 min | | 4 | 30 min | |
| 5 | 60 min | | 5 | 60 min | |
| 6 | Ladder | | 6 | Ladder | |
| 7 | 1 min | 4 µg dsRNA-Ache + 2 µL saliva (adults) | 7 | 1 min | 2 µg dsRNA-Ache + 2 µL water |
| 8 | 10 min | | 8 | 10 min | |
| 9 | 30 min | | 9 | 30 min | |
| 10 | 60 min | | 10 | 60 min | |
| 11 | Ladder | | 11 | Ladder | |
| 12 | 1 min | 2 µg dsRNA-Ache + 2 µg dsDNA-HhAGO2 + 2 µL saliva(adults) | 12 | 1 min | 2 µg dsRNA-Ache + 4 µg dsDNA-HhAGO2 + 2 µL saliva |
| 13 | 10 min | | 13 | 10 min | |
| 14 | 30 min | | 14 | 30 min | |
| 15 | 60 min | | 15 | 60 min | |
| 16 | | | 16 | | |
| 17 | | | 17 | | |

#### 4.1.2 Salivary gland extract

##### 4.1.2.1 Adult and 2nd instar nymphs

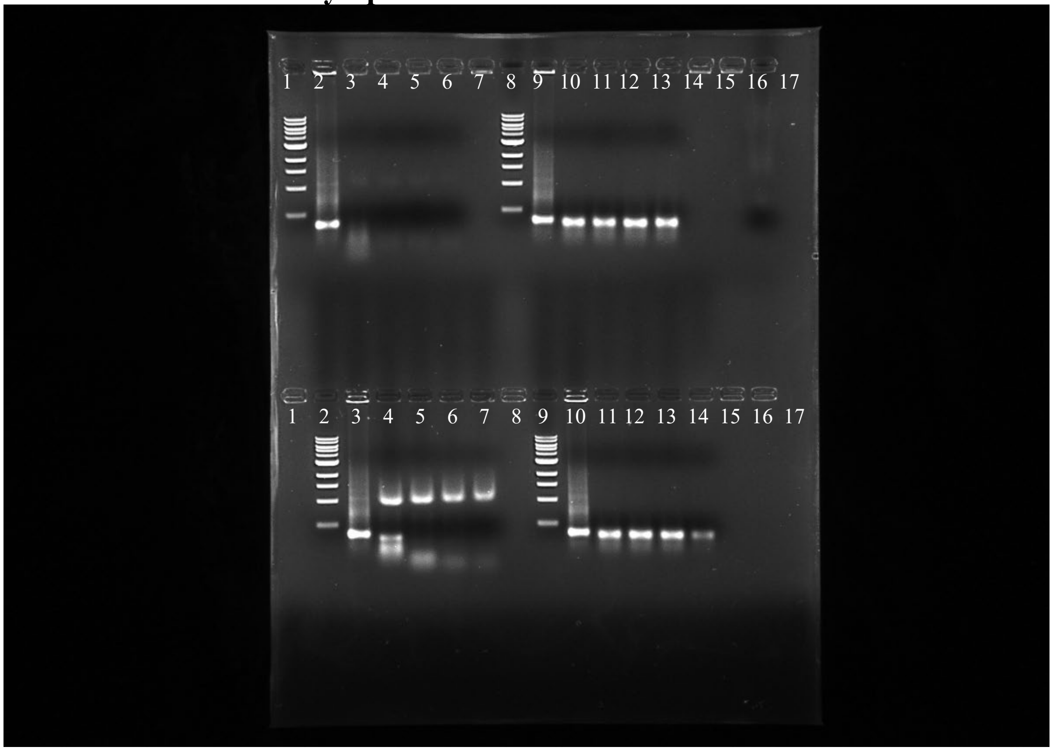

Consider upper lanes here: salivary gland extract

| well | sample name | Treatment | Well | Sample name | Treatment |
|---|---|---|---|---|---|
| 1 | Ladder | | 1 | | |
| 2 | dsRNA-GUS | | 2 | Ladder | |
| 3 | 1 min | | 3 | dsRNA-GUS | |
| 4 | 10 min | 2 µg dsRNA-GUS + 2 µL salivary gland extract (adults) | 4 | 1 min | 2 µg dsRNA-GUS + 2 µL salivary gland extract (2nd instar) |
| 5 | 30 min | | 5 | 10 min | |
| 6 | 60 min | | 6 | 30 min | |
| 7 | | | 7 | 60 min | |
| 8 | Ladder | | 8 | | |
| 9 | 1 min | | 9 | Ladder | |
| 10 | 10 min | | 10 | dsRNA-GUS | |
| 11 | 30 min | 2 µg dsRNA-GUS + 2 µL water | 11 | 1 min | 2 µg dsRNA-GUS + 2 µL water |
| 12 | 60 min | | 12 | 10 min | |
| 13 | | | 13 | 30 min | |
| 14 | | | 14 | 60 min | |
| 15 | | | 15 | | |
| 16 | | | 16 | | |
| 17 | | | 17 | | |

##### 4.1.2.2 5th stage

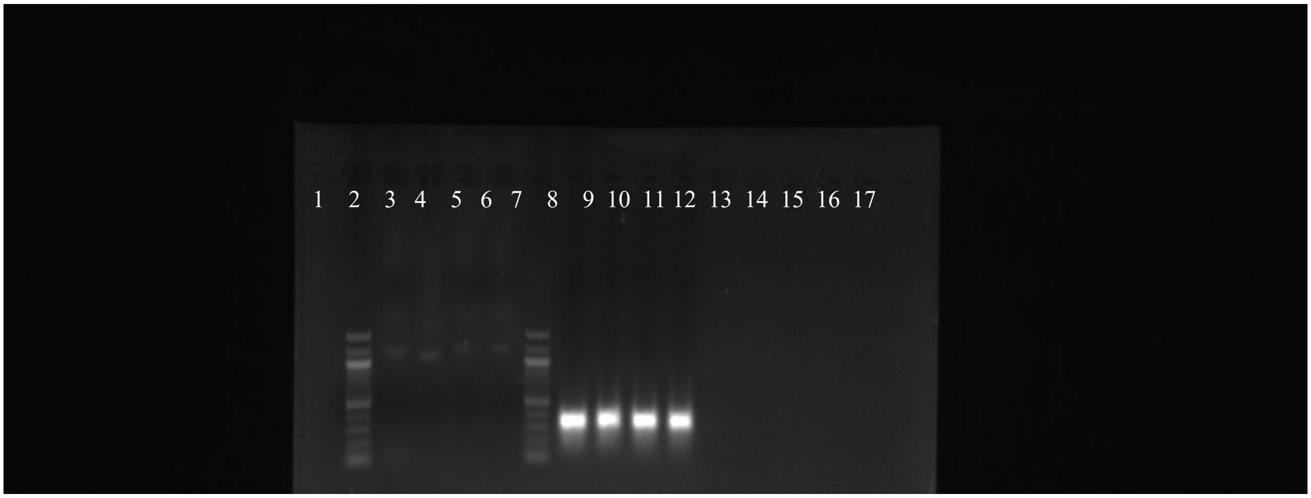

Above gel: Salivary gland extract 5th stage

| well | sample name | Treatment | Well | Sample name | Treatment |
|---|---|---|---|---|---|
| 1 | | | | | |
| 2 | Ladder | | | | |
| 3 | 1 min | 2 µg dsRNA-GUS + 2 µL salivary gland extract (5th stage) | | | |
| 4 | 10 min | | | | |
| 5 | 30 min | | | | |
| 6 | 60 min | | | | |
| 7 | Ladder | | | | |
| 8 | 1 min | 2 µg dsRNA-GUS + 2 µL water | | | |
| 9 | 10 min | | | | |
| 10 | 30 min | | | | |
| 11 | 60 min | | | | |
| 12 | | | | | |
| 13 | | | | | |
| 14 | | | | | |
| 15 | | | | | |
| 16 | | | | | |
| 17 | | | | | |

##### 4.1.2.3 $4^{th}$ stage

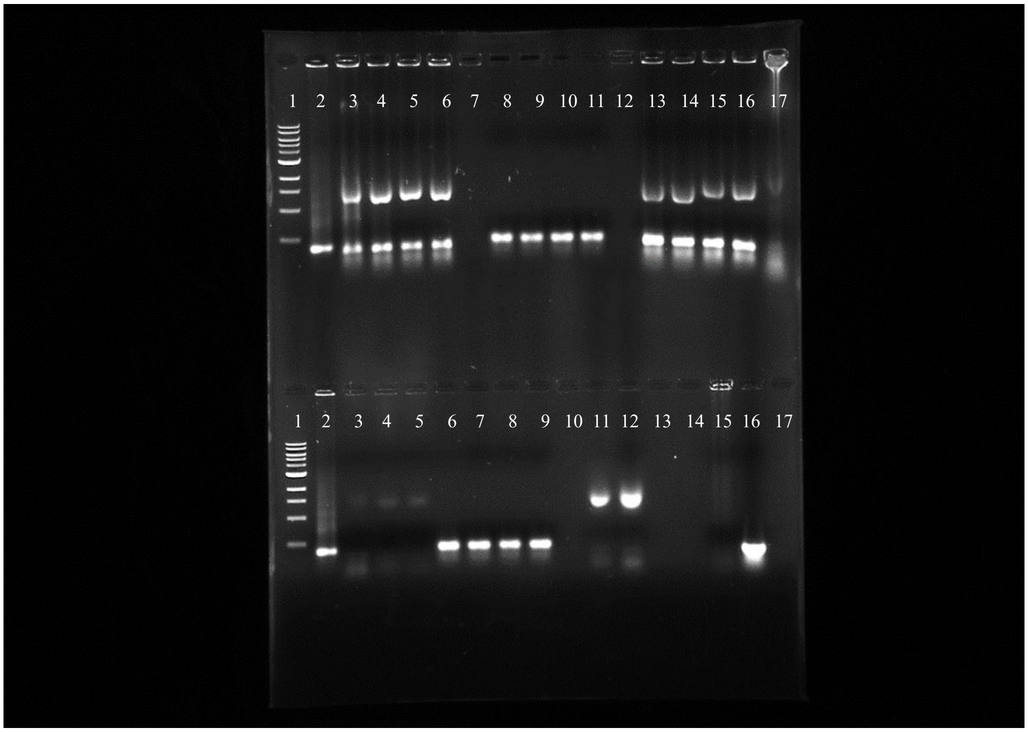

Consider only lower lanes: $4^{th}$ stage salivary gland extract10, 30, 60 min

| well | sample name | Treatment | Well | Sample name | Treatment |
|---|---|---|---|---|---|
| | | | 1 | Ladder | |
| | | | 2 | dsRNA-GUS | |
| | | | 3 | 10 min | 2 µg dsRNA-GUS + 2 µL salivary gland extract ($4^{th}$ instar) |
| | | | 4 | 30 min | |
| | | | 5 | 60 min | |
| | | | 6 | 1 min | 2 µg dsRNA-GUS + 2 µL water |
| | | | 7 | 10 min | |
| | | | 8 | 30 min | |
| | | | 9 | 60 min | |
| | | | 10 | | |
| | | | 11 | | |
| | | | 12 | | |
| | | | 13 | | |
| | | | 14 | | |
| | | | 15 | | |
| | | | 16 | | |
| | | | 17 | | |

#### 4.1.2.4 $3^{rd}$ stage

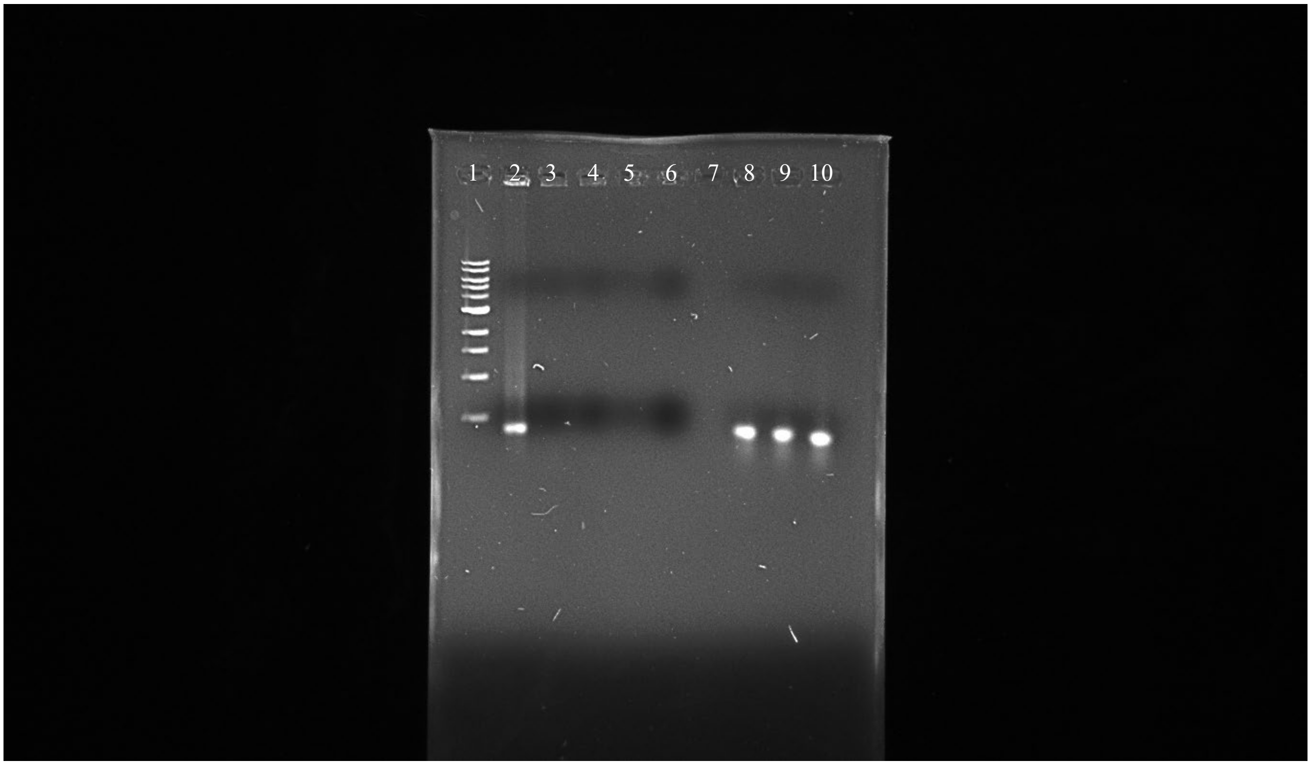

Above gel $3^{rd}$ stage Salivary gland extract- 03062022

| well | sample name | Treatment |
|---|---|---|
| 1 | Ladder | |
| 2 | dsRNA-GUS | |
| 3 | 1 min | 2 µg dsRNA-GUS + 2 µL salivary gland extract ($3^{rd}$ stage) |
| 4 | 10 min | |
| 5 | 30 min | |
| 6 | 60 min | |
| 7 | | |
| 8 | 1 min | 2 µg dsRNA-GUS + 2 µL water |
| 9 | 10 min | |
| 10 | 30 min | |

##### 4.1.2.5 Salivary gland extract : 2, 3-2, 4-2, 5-2 and A-2 stage

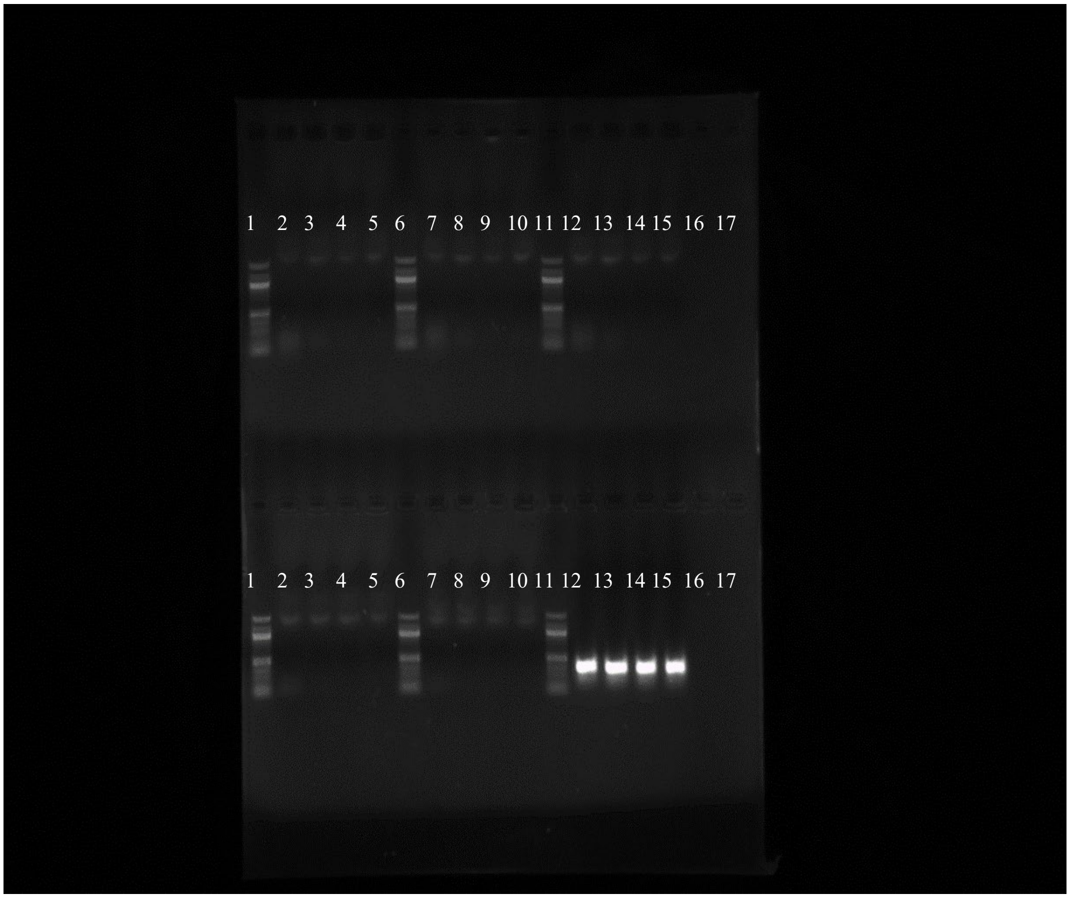

| well | sample name | Treatment | Well | Sample name | Treatment |
|---|---|---|---|---|---|
| 1 | Ladder | | 1 | Ladder | |
| 2 | 1 min | 2 µg dsRNA-GUS + 2 µL salivary gland extract (2nd instar) | 2 | 1 min | 2 µg dsRNA-GUS + 2 µL salivary gland extract (5th instar) |
| 3 | 10 min | | 3 | 10 min | |
| 4 | 30 min | | 4 | 30 min | |
| 5 | 60 min | | 5 | 60 min | |
| 6 | Ladder | | 6 | Ladder | |
| 7 | 1 min | 2 µg dsRNA-GUS + 2 µL salivary gland extract (3rd instar) | 7 | 1 min | 2 µg dsRNA-GUS + 2 µL salivary gland extract (adults) |
| 8 | 10 min | | 8 | 10 min | |
| 9 | 30 min | | 9 | 30 min | |
| 10 | 60 min | | 10 | 60 min | |
| 11 | Ladder | | 11 | Ladder | |
| 12 | 1 min | 2 µg dsRNA-GUS + 2 µL salivary gland extract (4th instar) | 12 | 1 min | 2 µg dsRNA-GUS + 2 µL water |
| 13 | 10 min | | 13 | 10 min | |
| 14 | 30 min | | 14 | 30 min | |
| 15 | 60 min | | 15 | 60 min | |
| 16 | | | 16 | | |
| 17 | | | 17 | | |

##### 4.1.3 Hemolymph

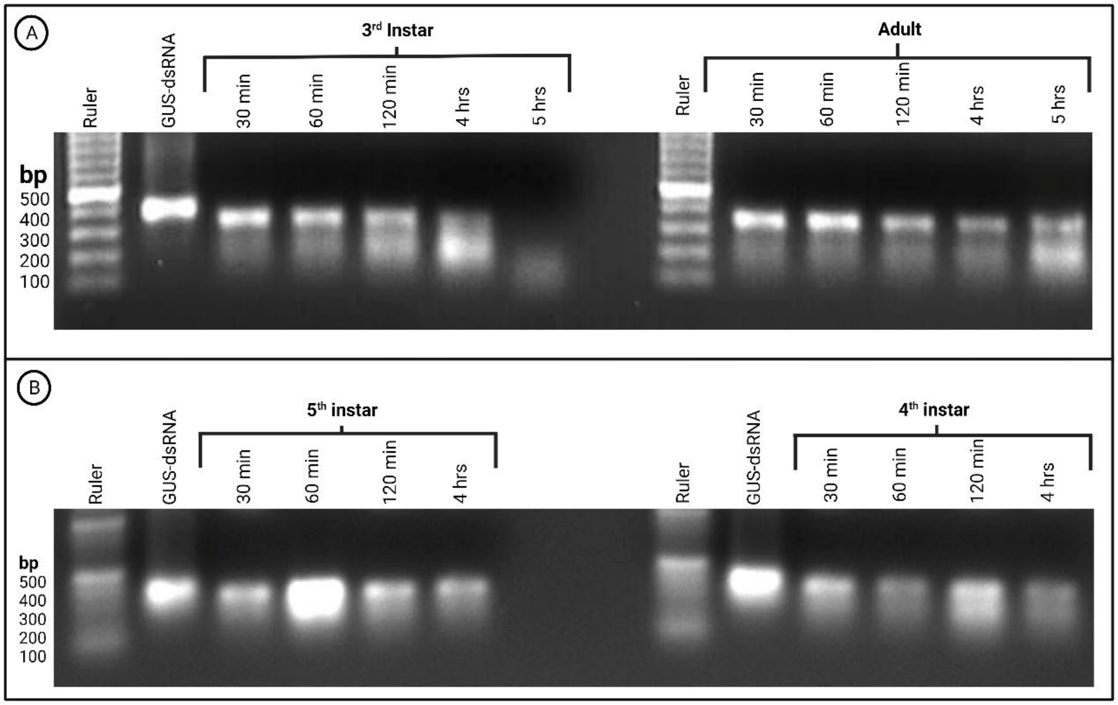

##### 4.1.3.1 3rd instar-1 and adult-1

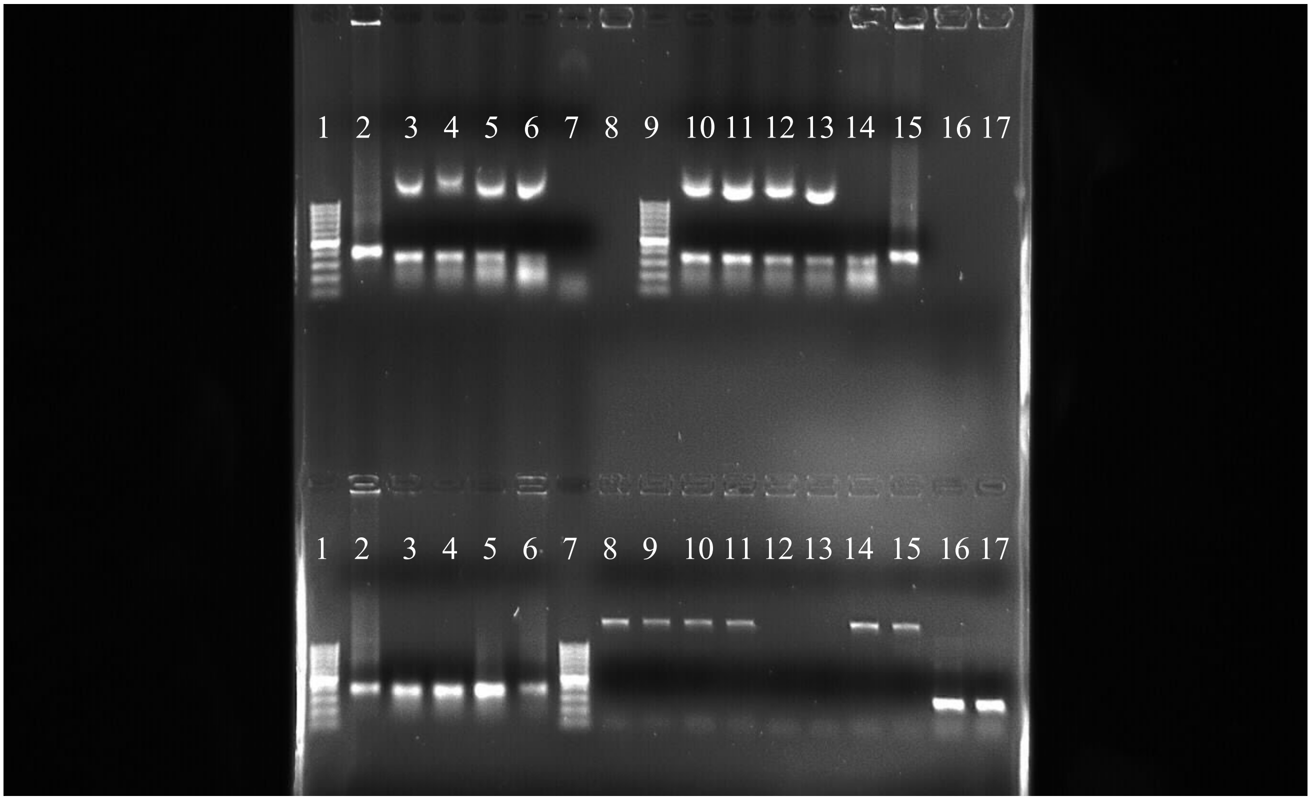

Above gel: 19082022

| well | sample name | Treatment | Well | Sample name | Treatment |
|---|---|---|---|---|---|
| 1 | Ladder | | 1 | Ladder | |
| 2 | dsRNA-GUS | | 2 | dsRNA-GUS | |
| 3 | 30 min | 2 µg dsRNA-GUS + 2 µL Hemolymph (3rd instar) | 3 | 30 min | 2 µg dsRNA-GUS + 2 µL water |
| 4 | 60 min | | 4 | 60 min | |
| 5 | 120 min | | 5 | 120 min | |
| 6 | 4 hrs. | | 6 | 4 hrs. | |
| 7 | 5 hrs. | | 7 | | |
| 8 | | | 8 | | |
| 9 | Ladder | | 9 | | |
| 10 | 30 min | 2 µg dsRNA-GUS + 2 µL Hemolymph (adults) | 10 | | |
| 11 | 60 min | | 11 | | |
| 12 | 120 min | | 12 | | |
| 13 | 4 hrs. | | 13 | | |
| 14 | 5 hrs. | | 14 | | |
| 15 | dsRNA-GUS | | 15 | | |
| 16 | | | 16 | | |
| 17 | | | 17 | | |

##### 4.1.3.2 5th instar-1 and 4th instar-1- 07 09 2022

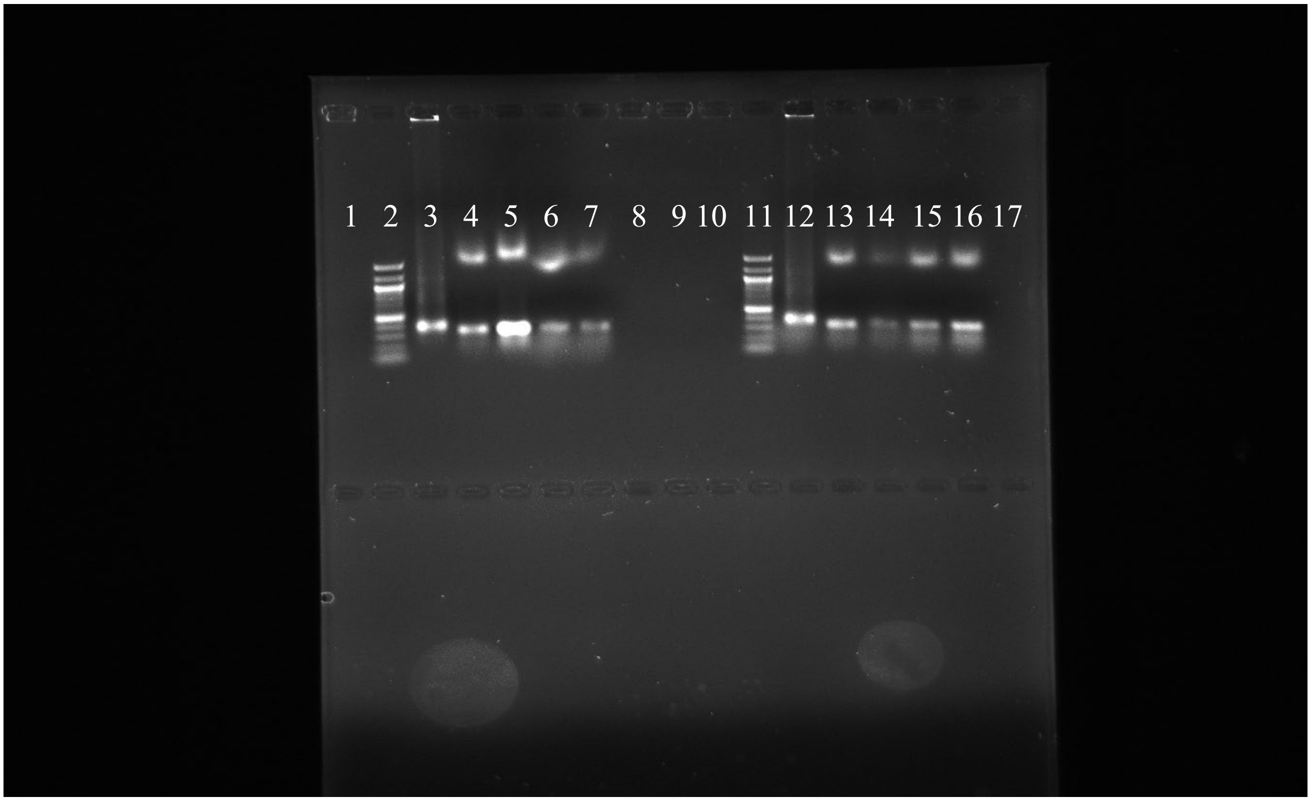

| well | sample name | Treatment | Well | Sample name | Treatment |
|---|---|---|---|---|---|
| 1 | | | 1 | | |
| 2 | Ladder | | 2 | | |
| 3 | dsRNA-GUS | | 3 | | |
| 4 | 30 min | | 4 | | |
| 5 | 60 min | 2 µg dsRNA-GUS + 2 µL Hemolymph (5th instar) | 5 | | |
| 6 | 120 min | | 6 | | |
| 7 | 4 hrs. | | 7 | | |
| 8 | | | 8 | | |
| 9 | | | 9 | | |
| 10 | | | 10 | | |
| 11 | Ladder | | 11 | | |
| 12 | dsRNA-GUS | | 12 | | |
| 13 | 30 min | | 13 | | |
| 14 | 60 min | 2 µg dsRNA-GUS + 2 µL Hemolymph (4th instar) | 14 | | |
| 15 | 120 min | | 15 | | |
| 16 | 4 hrs. | | 16 | | |
| 17 | | | 17 | | |

##### 4.1.3.3 Hemolymph - 2nd instar-1, 3rd instar-2, 4th instar-2

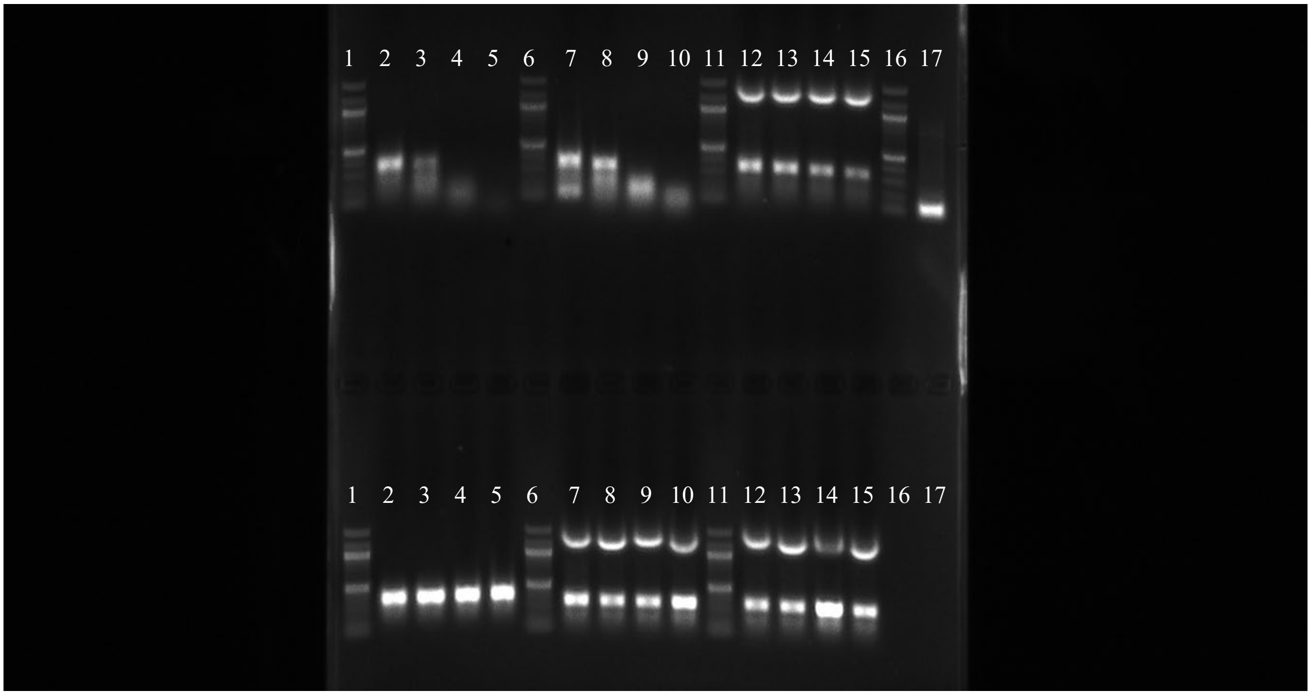

| well | sample name | Treatment | Well | Sample name | Treatment |
|---|---|---|---|---|---|
| 1 | | | 1 | Ladder | |
| 2 | | | 2 | 30 min | 2 µg dsRNA-GUS + 2 µL water |
| 3 | | | 3 | 60 min | |
| 4 | | | 4 | 120 min | |
| 5 | | | 5 | 4 hrs. | |
| 6 | | | 6 | Ladder | |
| 7 | | | 7 | 30 min | 2 µg dsRNA-GUS + 2 µL Hemolymph (3rd instar) |
| 8 | | | 8 | 60 min | |
| 9 | | | 9 | 120 min | |
| 10 | | | 10 | 4 hrs. | |
| 11 | Ladder | | 11 | Ladder | |
| 12 | 30 min | 2 µg dsRNA-GUS + 2 µL Hemolymph (2nd instar) | 12 | 30 min | 2 µg dsRNA-GUS + 2 µL Hemolymph (4th instar) |
| 13 | 60 min | | 13 | 60 min | |
| 14 | 120 min | | 14 | 120 min | |
| 15 | 4 hrs. | | 15 | 4 hrs. | |
| 16 | | | 16 | | |
| 17 | | | 17 | | |

##### 4.1.3.4 Hemolymph - Adult-3, 5$^{th}$ instar-2, 4$^{th}$ instar-3

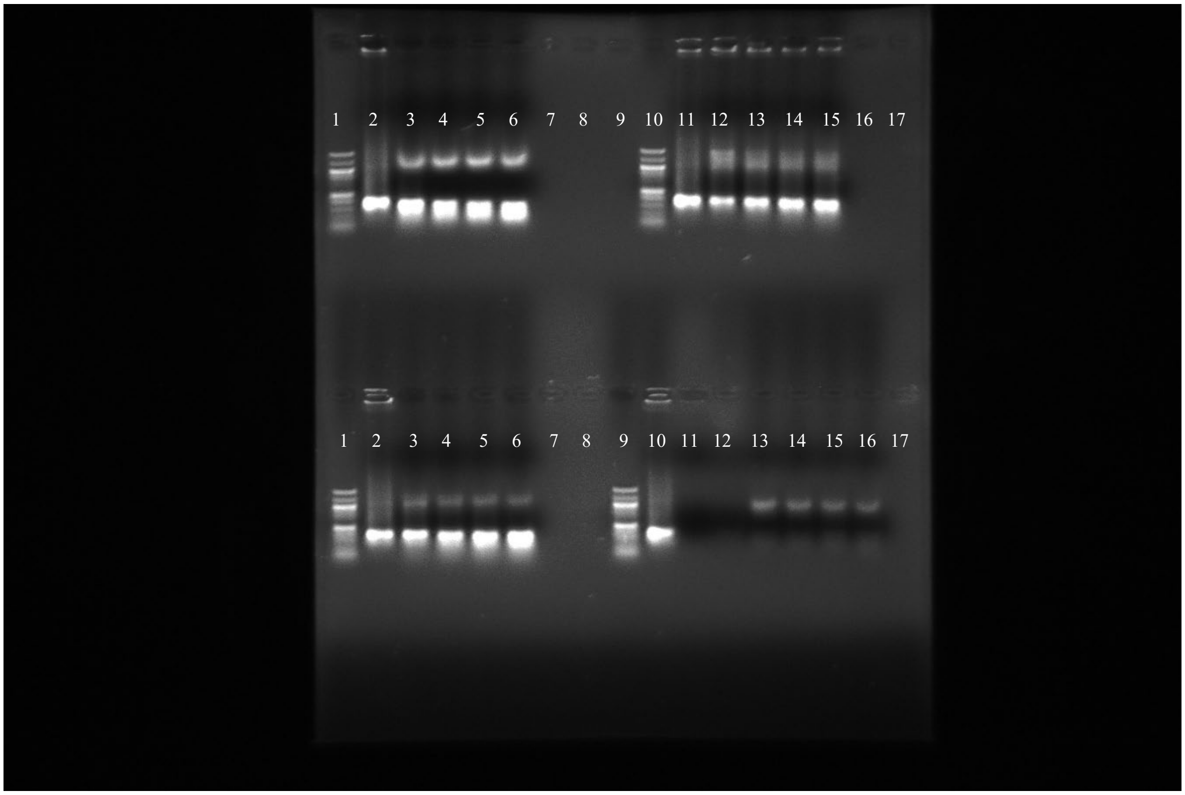

Above gel: 24062022

| Well (upper) | sample | Treatment | Well (Lower) | Sample | Treatment |
|---|---|---|---|---|---|
| 1 | Ladder | | 1 | Ladder | |
| 2 | dsRNA-GUS | | 2 | dsRNA-GUS | |
| 3 | 30 min | | 3 | 30 min | 2 µg dsRNA-GUS + 2 µL Hemolymph (4$^{th}$ instar) |
| 4 | 60 min | 2 µg dsRNA-GUS + 2 µL Hemolymph (adults) | 4 | 60 min | |
| 5 | 120 min | | 5 | 120 min | |
| 6 | 4 hrs. | | 6 | 4 hrs. | |
| 7 | | | 7 | | |
| 8 | | | 8 | | |
| 9 | | | 9 | | |
| 10 | Ladder | | 10 | | |
| 11 | dsRNA-GUS | | 11 | | |
| 12 | 30 min | | 12 | | |
| 13 | 60 min | | 13 | | |
| 14 | 120 min | 2 µg dsRNA-GUS + 2 µL Hemolymph (5$^{th}$ instar) | 14 | | |
| 15 | 4 hrs. | | 15 | | |
| 16 | | | 16 | | |
| 17 | | | 17 | | |

#### 4.1.3.5 Hemolymph – 3rd instar-3, 2nd instar-2

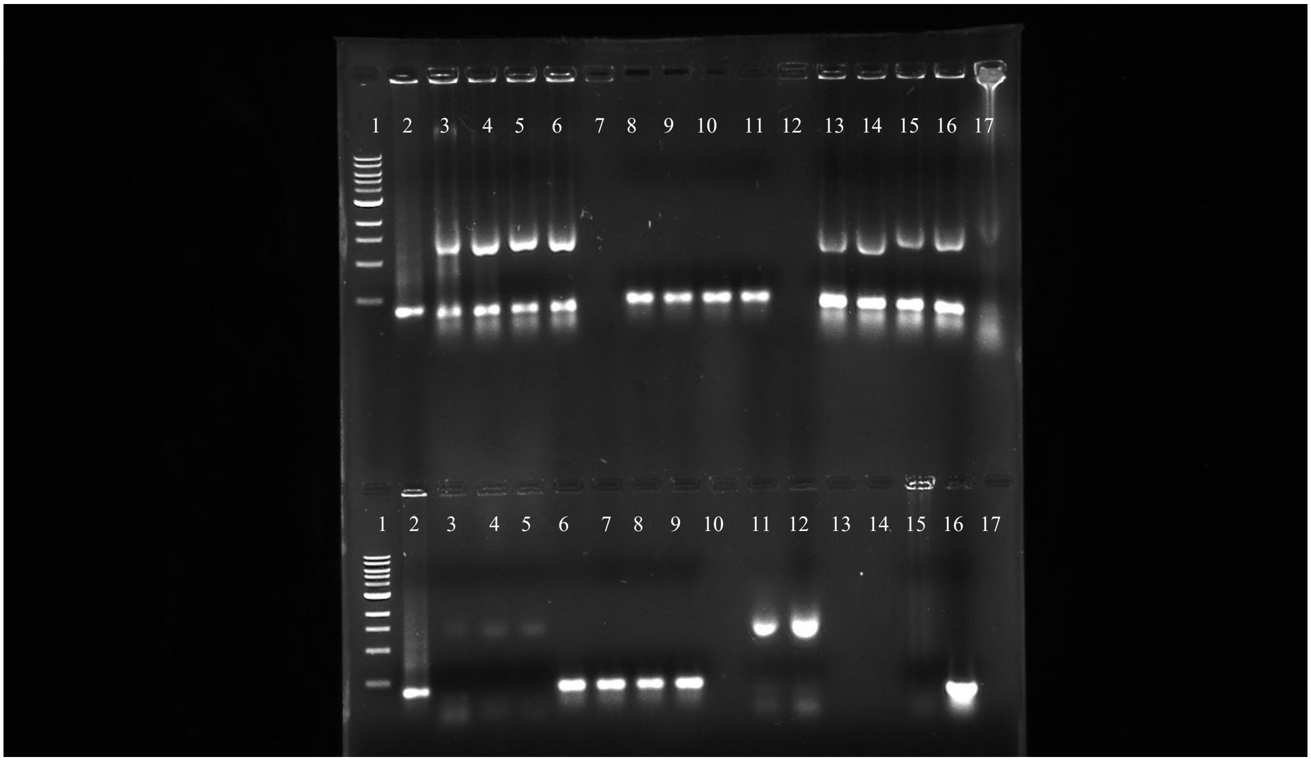

| Well (upper) | sample | Treatment | Well (Lower) | Sample | Treatment |
|---|---|---|---|---|---|
| 1 | Ladder | | 1 | | |
| 2 | dsRNA-GUS | | 2 | | |
| 3 | 30 min | 2 µg dsRNA-GUS + 2 µL Hemolymph (3rd instar) | 3 | | |
| 4 | 60 min | | 4 | | |
| 5 | 120 min | | 5 | | |
| 6 | 4 hrs. | | 6 | | |
| 7 | | | 7 | | |
| 8 | 30 min | 2 µg dsRNA-GUS + 2 µL water | 8 | | |
| 9 | 60 min | | 9 | | |
| 10 | 120 min | | 10 | | |
| 11 | 4 hrs. | | 11 | | |
| 12 | | | 12 | | |
| 13 | 30 min | 2 µg dsRNA-GUS + 2 µL Hemolymph (2nd instar) | 13 | | |
| 14 | 60 min | | 14 | | |
| 15 | 120 min | | 15 | | |
| 16 | 4 hrs. | | 16 | | |
| 17 | | | 17 | | |

#### 5 Statistical analysis of dsRNA stability assays in saliva, salivary gland extract and hemolymph of *H. halys*:

In the present study, a comprehensive statistical analysis was performed using SAS software (version 9.4) to evaluate the effect of treatment, life stage, and their interaction on haemolymph volume in experimental subjects. The data set was analysed using the PROC MIXED procedure and a linear mixed effects model. The fixed effects included treatment, life stage and their interaction, with time as a covariate. The model accounted for inherent variability by including random effects for gels and the replicate by gel interaction. The assumption of compound symmetry was accounted for by specifying replication, and the degrees of freedom method was set to Kenward-Roger. Post-hoc pairwise comparisons of treatment means were performed using the LSMEANS statement with the PDIFF option. The results provide critical insight into the specific effects of treatments on band volume on the agarose gel (dsRNA stability), taking into account the variability introduced by gels and replicates. This statistical approach enhances the reliability and interpretability of the results, thus contributing to the robustness of the overall experimental conclusions.

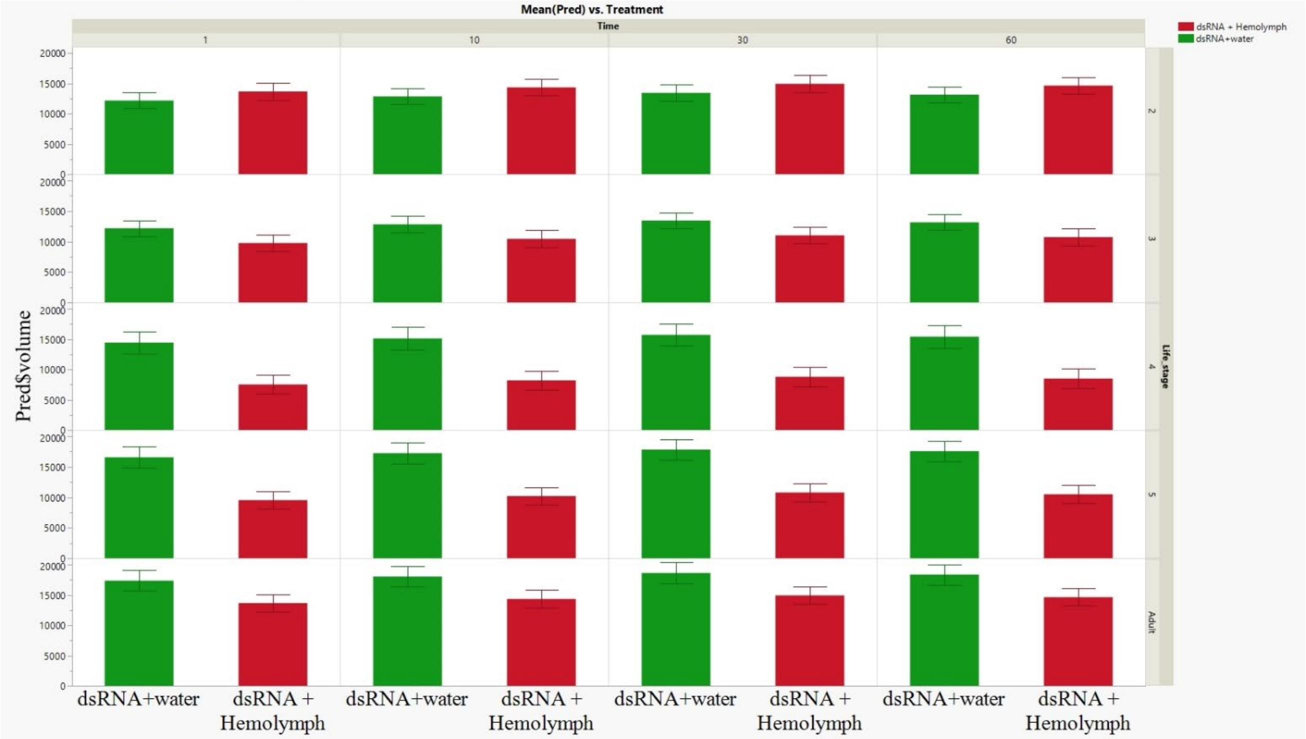

Supplementary figure 4. Stability of dsRNA when subjected to ex-vivo incubation with *H. halys* hemolymph. The Y-axis represents the predicted volume of the dsRNA band on the gel image captured under UV light, while the X-axis denotes various treatments employed during dsRNA incubation. The columns represent incubation time, the rows represent different life stages (2nd, 3rd, 4th, 5th instar, and adults) during which dsRNA stability was evaluated. The experiments are replicated 2 – 3 times for robustness. The gel images were processed using ImageJ software. Statistical analysis was executed using SAS, and the graphical representation was generated using JMP-Pro. This figure provides insights into the differential stability of dsRNA in hemolymph across different life stages, contributing valuable information to the experimental outcomes.

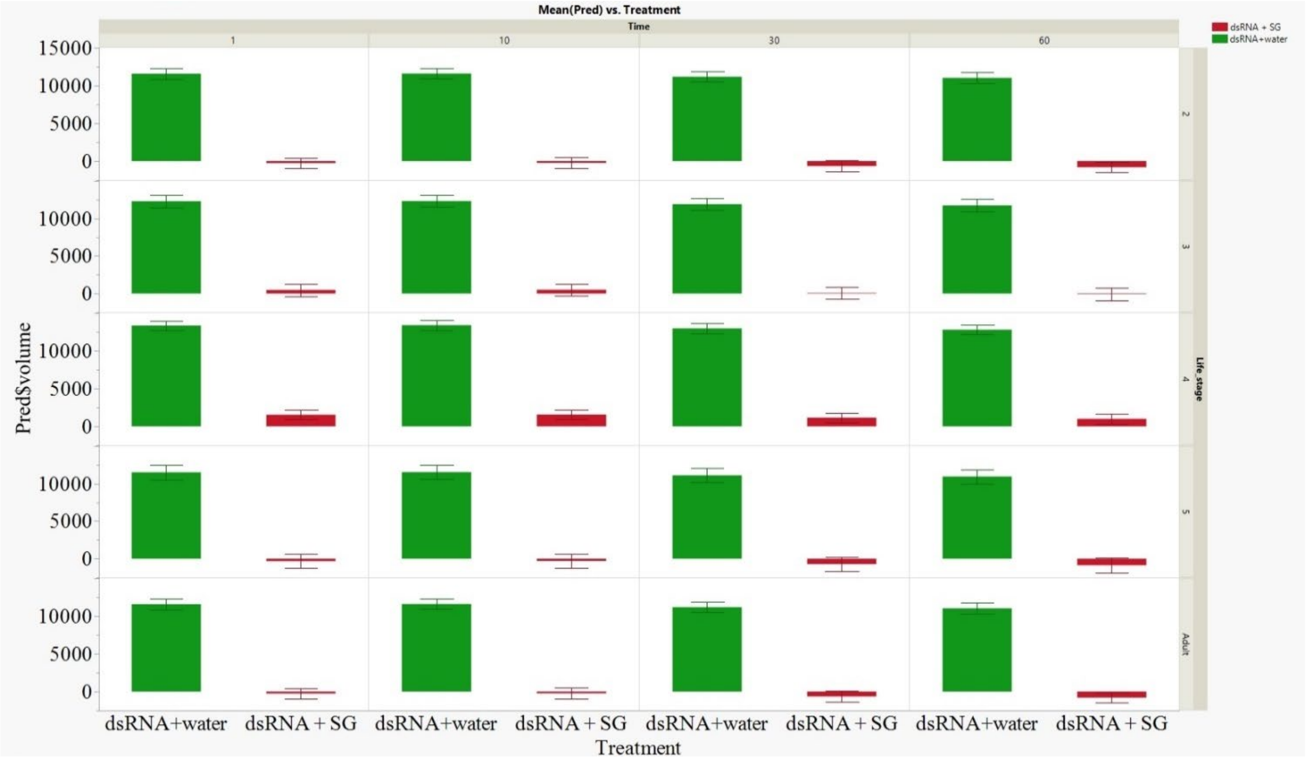

Supplementary figure 5. Stability of dsRNA when subjected to ex-vivo incubation with *H. halys* salivary gland extract. The Y-axis represents the predicted volume of the dsRNA band on the gel image captured under UV light, while the X-axis denotes various treatments employed during dsRNA incubation. Additionally, the Y-axis incorporates an overlay denoting different life stages (2nd, 3rd, 4th, 5th instar, and adults) during which dsRNA stability was evaluated. The experiments were replicated twice for robustness. The gel images are processed using ImageJ software. Statistical analysis was executed using SAS, and the graphical representation was generated using JMP-Pro. This figure provides insights into the differential stability of dsRNA in salivary gland extract across different life stages, contributing valuable information to the experimental outcomes.

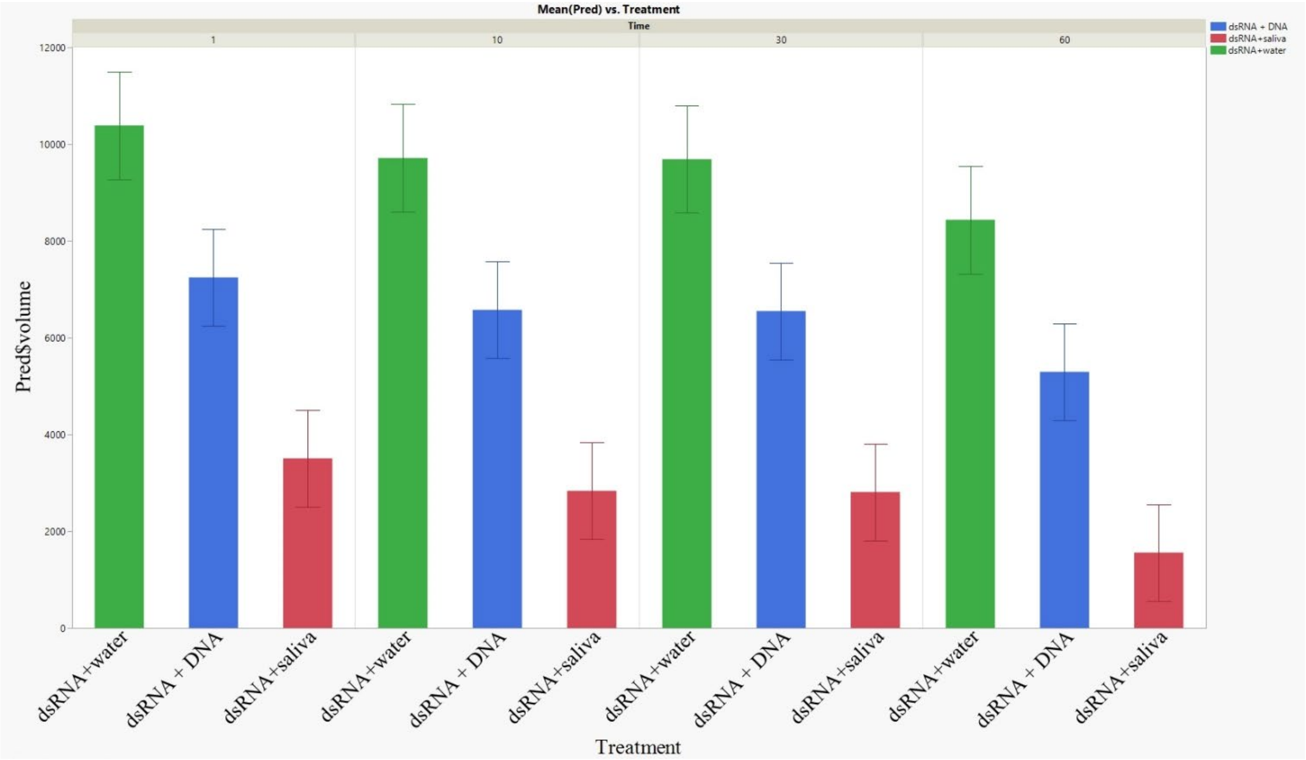

Supplementary figure 6 Stability of dsRNA when subjected to ex-vivo incubation with *H. halys* watery saliva collected from tip of its rostrum/proboscis. The Y-axis represents the predicted volume of the dsRNA band on the gel image captured under UV light, while the X-axis denotes various treatments employed during dsRNA incubation. One of the three different treatments involve dsRNA incubation with dsDNA at a 1:1 ratio. The experiments were replicated thrice for robustness. The gel images are processed using ImageJ software. Statistical analysis was executed using SAS, and the graphical representation was generated using JMP-Pro. This figure provides insights into the differential stability of dsRNA in salivary gland extract across different life stages, contributing valuable information to the experimental outcomes.
