## Supplementary material for "Double-stranded DNA reduces dsRNA degradation in the saliva and significantly enhanced RNAi-mediated gene silencing in *Halyomorpha halys*": ex_vivo_data

### The SAS System

#### The Mixed Procedure

| Model Information | |
|---|---|
| Data Set | WORK.GEL_DATA |
| Dependent Variable | Volume |
| Covariance Structure | Variance Components |
| Estimation Method | REML |
| Residual Variance Method | Profile |
| Fixed Effects SE Method | Kenward-Roger |
| Degrees of Freedom Method | Kenward-Roger |

| Class Level Information | | |
|---|---|---|
| Class | Levels | Values |
| Treatment | 2 | dsRNA + Hemolymph dsRNA+water |
| Time | 5 | 0 1 10 30 60 |
| Gels | 4 | 6 9 10 11 |
| Life_stage | 5 | 2 3 4 5 Adult |
| Repetition | 1 | A |

| Dimensions | |
|---|---|
| Covariance Parameters | 3 |
| Columns in X | 23 |
| Columns in Z | 8 |
| Subjects | 1 |
| Max Obs per Subject | 78 |

| Number of Observations | |
|---|---|
| Number of Observations Read | 90 |
| Number of Observations Used | 78 |
| Number of Observations Not Used | 12 |

| Iteration History | | | |
|---|---|---|---|
| Iteration | Evaluations | -2 Res Log Like | Criterion |
| 0 | 1 | 1277.62911482 | |
| 1 | 2 | 1249.67564415 | 0.00169842 |
| 2 | 1 | 1249.46555118 | 0.00017579 |
| 3 | 1 | 1249.42060929 | 0.00000813 |
| 4 | 1 | 1249.41748012 | 0.00000005 |
| 5 | 1 | 1249.41745803 | 0.00000000 |

Convergence criteria met but final Hessian is not positive definite.

| Covariance Parameter Estimates | |
|---|---|
| **Cov Parm** | **Estimate** |
| **Gels** | 13551123 |
| **Gels*Repetition** | 4940367 |
| **Residual** | 9734101 |

| Fit Statistics | |
|---|---|
| **-2 Res Log Likelihood** | 1249.4 |
| **AIC (Smaller is Better)** | 1255.4 |
| **AICC (Smaller is Better)** | 1255.8 |
| **BIC (Smaller is Better)** | 1253.6 |

**Solution for Fixed Effects**

| Effect | Treatment | Life_stage | Time | Estimate | Standard Error | DF | t Value | Pr > \|t\| |
|---|---|---|---|---|---|---|---|---|
| **Intercept** | | | | 19940 | 2898.90 | 6.79 | 6.88 | 0.0003 |
| **Treatment** | dsRNA + Hemolymph | | | -3709.75 | 1811.65 | 61.5 | -2.05 | 0.0449 |
| **Treatment** | dsRNA+water | | | 0 | . | . | . | . |
| **Life_stage** | | 2 | | -8376.43 | 2761.79 | 56.7 | -3.03 | 0.0036 |
| **Life_stage** | | 3 | | -8376.43 | 2761.79 | 56.7 | -3.03 | 0.0036 |
| **Life_stage** | | 4 | | -6371.50 | 2628.67 | 61.7 | -2.42 | 0.0183 |
| **Life_stage** | | 5 | | -787.70 | 1801.31 | 60.9 | -0.44 | 0.6634 |
| **Life_stage** | | Adult | | 0 | . | . | . | . |
| **Treatment*Life_stage** | dsRNA + Hemolymph | 2 | | 5202.65 | 2300.13 | 61.2 | 2.26 | 0.0273 |
| **Treatment*Life_stage** | dsRNA + Hemolymph | 3 | | 1301.46 | 2300.13 | 61.2 | 0.57 | 0.5736 |
| **Treatment*Life_stage** | dsRNA + Hemolymph | 4 | | -3204.04 | 2502.93 | 62.5 | -1.28 | 0.2052 |
| **Treatment*Life_stage** | dsRNA + Hemolymph | 5 | | -3343.09 | 2382.90 | 60.9 | -1.40 | 0.1657 |
| **Treatment*Life_stage** | dsRNA + Hemolymph | Adult | | 0 | . | . | . | . |
| **Treatment*Life_stage** | dsRNA+water | 2 | | 0 | . | . | . | . |
| **Treatment*Life_stage** | dsRNA+water | 3 | | 0 | . | . | . | . |
| **Treatment*Life_stage** | dsRNA+water | 4 | | 0 | . | . | . | . |
| **Treatment*Life_stage** | dsRNA+water | 5 | | 0 | . | . | . | . |
| **Treatment*Life_stage** | dsRNA+water | Adult | | 0 | . | . | . | . |
| **Time** | | | 0 | -3225.22 | 1370.46 | 61.2 | -2.35 | 0.0218 |
| **Time** | | | 1 | -978.42 | 1070.13 | 60.9 | -0.91 | 0.3642 |
| **Time** | | | 10 | -308.03 | 1070.13 | 60.9 | -0.29 | 0.7744 |
| **Time** | | | 30 | 285.05 | 1070.13 | 60.9 | 0.27 | 0.7909 |
| **Time** | | | 60 | 0 | . | . | . | . |

| Type 3 Tests of Fixed Effects | | | | |
|---|---|---|---|---|
| **Effect** | **Num DF** | **Den DF** | **F Value** | **Pr > F** |
| **Treatment** | 1 | 61.7 | 19.24 | <.0001 |
| **Life_stage** | 4 | 61 | 5.54 | 0.0007 |
| **Treatment*Life_stage** | 4 | 61.5 | 4.78 | 0.0020 |
| **Time** | 4 | 61 | 1.91 | 0.1198 |

| Least Squares Means | | | | | | |
|---|---|---|---|---|---|---|
| **Effect** | **Treatment** | **Estimate** | **Standard Error** | **DF** | **t Value** | **Pr > \|t\|** |
| **Treatment** | dsRNA + Hemolymph | 10594 | 2222.28 | 2.79 | 4.77 | 0.0206 |
| **Treatment** | dsRNA+water | 14312 | 2223.83 | 2.8 | 6.44 | 0.0094 |

| Differences of Least Squares Means | | | | | | | |
|---|---|---|---|---|---|---|---|
| **Effect** | **Treatment** | **Treatment** | **Estimate** | **Standard Error** | **DF** | **t Value** | **Pr > \|t\|** |
| **Treatment** | dsRNA + Hemolymph | dsRNA+water | -3718.35 | 847.74 | 61.7 | -4.39 | <.0001 |

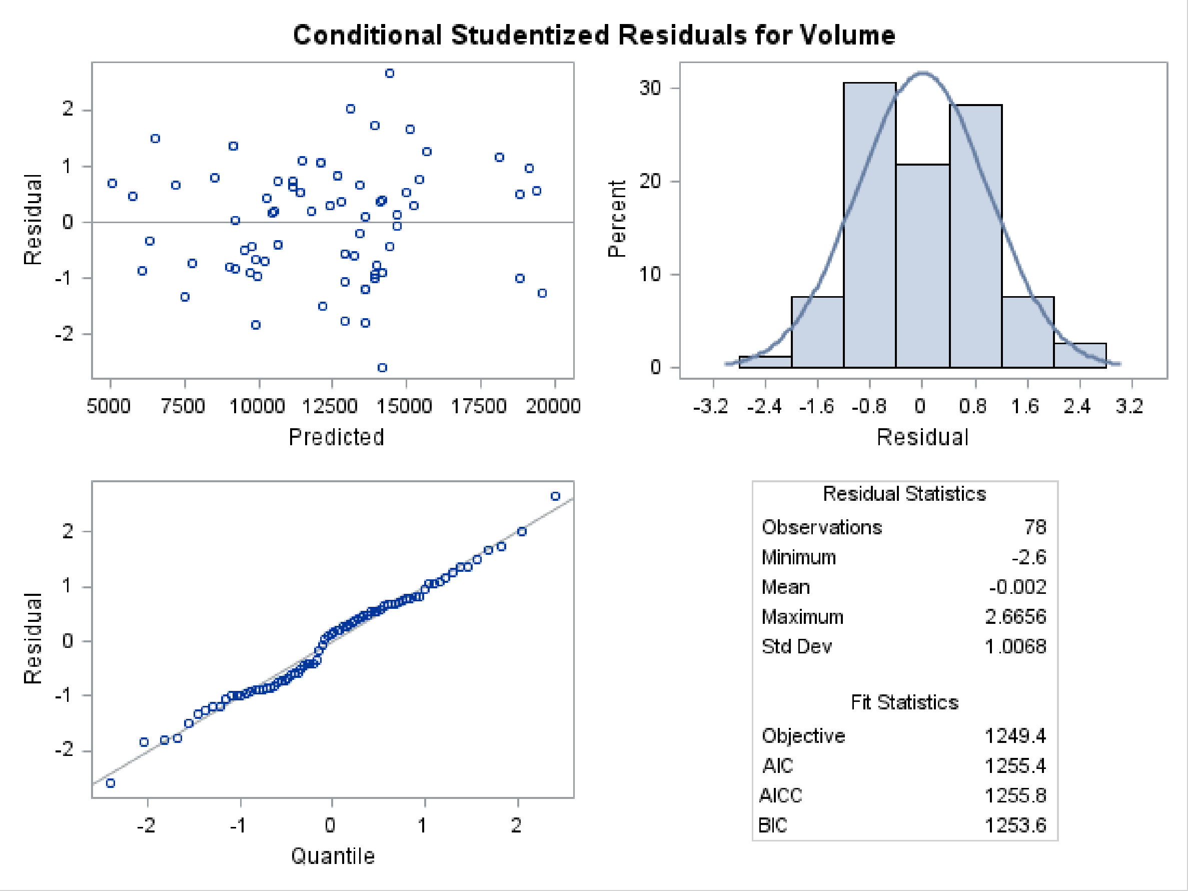

### The SAS System

#### The Mixed Procedure

| Model Information | |
|---|---|
| Data Set | WORK.GEL_DATA |
| Dependent Variable | Volume |
| Covariance Structure | Variance Components |
| Estimation Method | REML |
| Residual Variance Method | Profile |
| Fixed Effects SE Method | Kenward-Roger |
| Degrees of Freedom Method | Kenward-Roger |

| Class Level Information | | |
|---|---|---|
| Class | Levels | Values |
| Treatment | 3 | dsRNA + DNA dsRNA+saliva dsRNA+water |
| Time | 5 | 0 1 10 30 60 |
| Gels | 3 | 1 2 3 |
| Life_stage | 1 | Adult |
| Repetition | 1 | A |

| Dimensions | |
|---|---|
| Covariance Parameters | 2 |
| Columns in X | 9 |
| Columns in Z | 3 |
| Subjects | 1 |
| Max Obs per Subject | 34 |

| Number of Observations | |
|---|---|
| Number of Observations Read | 39 |
| Number of Observations Used | 34 |
| Number of Observations Not Used | 5 |

| Iteration History | | | |
|---|---|---|---|
| Iteration | Evaluations | -2 Res Log Like | Criterion |
| 0 | 1 | 522.86979103 | |
| 1 | 2 | 506.06037367 | 0.00002689 |
| 2 | 1 | 506.05370833 | 0.00000056 |
| 3 | 1 | 506.05357818 | 0.00000000 |

Convergence criteria met.

| Covariance Parameter Estimates |
|---|

| Cov Parm | Estimate |
|---|---|
| Gels | 7460110 |
| Residual | 4058731 |

| Fit Statistics | |
|---|---|
| -2 Res Log Likelihood | 506.1 |
| AIC (Smaller is Better) | 510.1 |
| AICC (Smaller is Better) | 510.6 |
| BIC (Smaller is Better) | 508.3 |

**Solution for Fixed Effects**

| Effect | Treatment | Time | Estimate | Standard Error | DF | t Value | Pr > \|t\| |
|---|---|---|---|---|---|---|---|
| Intercept | | | 8435.62 | 1859.35 | 3.4 | 4.54 | 0.0152 |
| Treatment | dsRNA + DNA | | -3139.04 | 963.43 | 25.2 | -3.26 | 0.0032 |
| Treatment | dsRNA+saliva | | -6878.57 | 963.43 | 25.2 | -7.14 | <.0001 |
| Treatment | dsRNA+water | | 0 | . | . | . | . |
| Time | | 0 | 2497.40 | 1776.12 | 25.2 | 1.41 | 0.1719 |
| Time | | 1 | 1950.69 | 1007.31 | 25 | 1.94 | 0.0642 |
| Time | | 10 | 1276.76 | 1007.31 | 25 | 1.27 | 0.2167 |
| Time | | 30 | 1255.29 | 1007.31 | 25 | 1.25 | 0.2243 |
| Time | | 60 | 0 | . | . | . | . |

**Type 3 Tests of Fixed Effects**

| Effect | Num DF | Den DF | F Value | Pr > F |
|---|---|---|---|---|
| Treatment | 2 | 25.1 | 26.86 | <.0001 |
| Time | 4 | 25 | 1.15 | 0.3562 |

**Least Squares Means**

| Effect | Treatment | Estimate | Standard Error | DF | t Value | Pr > \|t\| |
|---|---|---|---|---|---|---|
| Treatment | dsRNA + DNA | 6692.60 | 1713.45 | 2.49 | 3.91 | 0.0414 |
| Treatment | dsRNA+saliva | 2953.07 | 1713.45 | 2.49 | 1.72 | 0.2017 |
| Treatment | dsRNA+water | 9831.64 | 1712.91 | 2.48 | 5.74 | 0.0174 |

**Differences of Least Squares Means**

| Effect | Treatment | Treatment | Estimate | Standard Error | DF | t Value | Pr > \|t\| | Adjustment | Adj P |
|---|---|---|---|---|---|---|---|---|---|
| Treatment | dsRNA + DNA | dsRNA+saliva | 3739.53 | 822.47 | 25 | 4.55 | 0.0001 | Tukey-Kramer | 0.0003 |
| Treatment | dsRNA + DNA | dsRNA+water | -3139.04 | 963.43 | 25.2 | -3.26 | 0.0032 | Tukey-Kramer | 0.0087 |
| Treatment | dsRNA+saliva | dsRNA+water | -6878.57 | 963.43 | 25.2 | -7.14 | <.0001 | Tukey-Kramer | <.0001 |

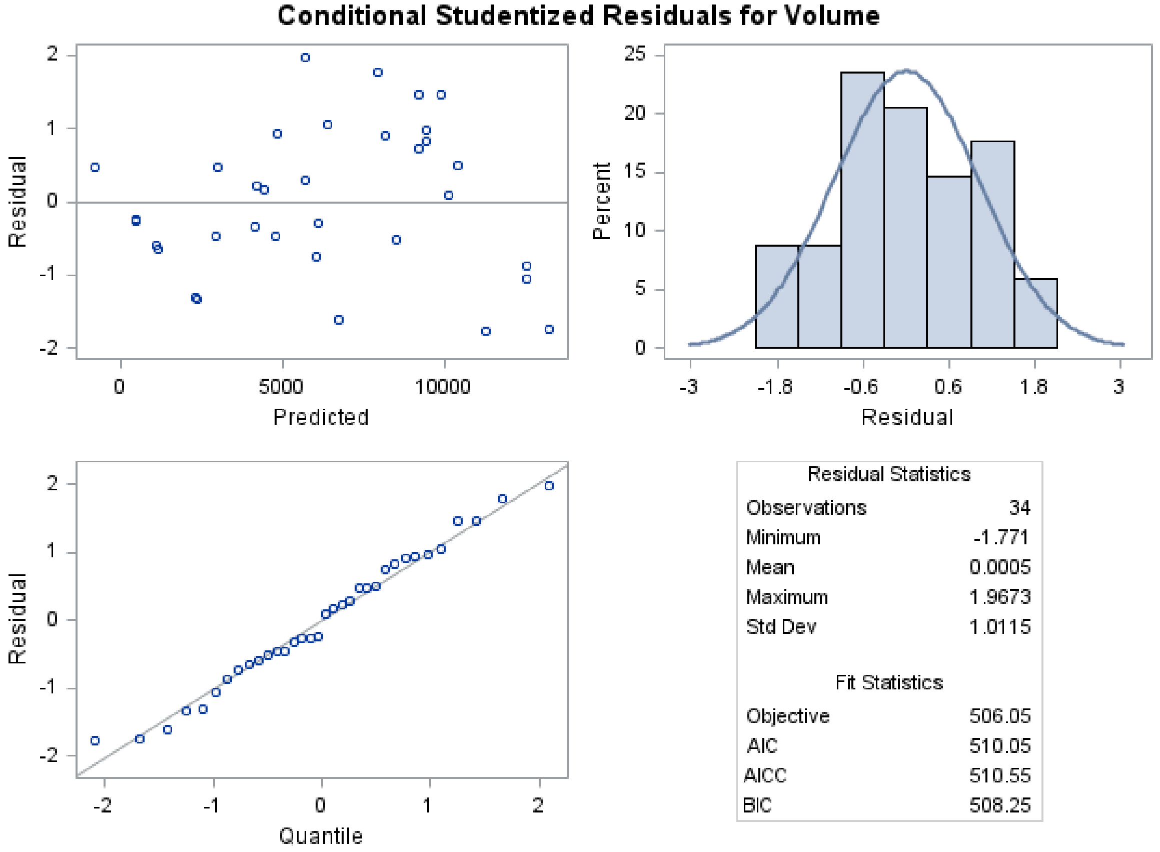

| Residual Statistics | |
|---|---|
| Observations | 34 |
| Minimum | -1.771 |
| Mean | 0.0005 |
| Maximum | 1.9673 |
| Std Dev | 1.0115 |

| Fit Statistics | |
|---|---|
| Objective | 506.05 |
| AIC | 510.05 |
| AICC | 510.55 |
| BIC | 508.25 |

### The SAS System

#### The Mixed Procedure

| Model Information | |
|---|---|
| Data Set | WORK.GEL_DATA |
| Dependent Variable | Volume |
| Covariance Structure | Variance Components |
| Estimation Method | REML |
| Residual Variance Method | Profile |
| Fixed Effects SE Method | Kenward-Roger |
| Degrees of Freedom Method | Kenward-Roger |

| Class Level Information | | |
|---|---|---|
| Class | Levels | Values |
| Treatment | 3 | dsRNA + DNA dsRNA+saliva dsRNA+water |
| Time | 5 | 0 1 10 30 60 |
| Gels | 3 | 1 2 3 |
| Life_stage | 1 | Adult |
| Repetition | 1 | A |

| Dimensions | |
|---|---|
| Covariance Parameters | 2 |
| Columns in X | 9 |
| Columns in Z | 3 |
| Subjects | 1 |
| Max Obs per Subject | 34 |

| Number of Observations | |
|---|---|
| Number of Observations Read | 39 |
| Number of Observations Used | 34 |
| Number of Observations Not Used | 5 |

| Iteration History | | | |
|---|---|---|---|
| Iteration | Evaluations | -2 Res Log Like | Criterion |
| 0 | 1 | 522.86979103 | |
| 1 | 2 | 506.06037367 | 0.00002689 |
| 2 | 1 | 506.05370833 | 0.00000056 |
| 3 | 1 | 506.05357818 | 0.00000000 |

Convergence criteria met.

| Covariance Parameter Estimates |
|---|

| Cov Parm | Estimate |
|---|---|
| Gels | 7460110 |
| Residual | 4058731 |

| Fit Statistics | |
|---|---|
| -2 Res Log Likelihood | 506.1 |
| AIC (Smaller is Better) | 510.1 |
| AICC (Smaller is Better) | 510.6 |
| BIC (Smaller is Better) | 508.3 |

**Solution for Fixed Effects**

| Effect | Treatment | Time | Estimate | Standard Error | DF | t Value | Pr > \|t\| |
|---|---|---|---|---|---|---|---|
| Intercept | | | 8435.62 | 1859.35 | 3.4 | 4.54 | 0.0152 |
| Treatment | dsRNA + DNA | | -3139.04 | 963.43 | 25.2 | -3.26 | 0.0032 |
| Treatment | dsRNA+saliva | | -6878.57 | 963.43 | 25.2 | -7.14 | <.0001 |
| Treatment | dsRNA+water | | 0 | . | . | . | . |
| Time | | 0 | 2497.40 | 1776.12 | 25.2 | 1.41 | 0.1719 |
| Time | | 1 | 1950.69 | 1007.31 | 25 | 1.94 | 0.0642 |
| Time | | 10 | 1276.76 | 1007.31 | 25 | 1.27 | 0.2167 |
| Time | | 30 | 1255.29 | 1007.31 | 25 | 1.25 | 0.2243 |
| Time | | 60 | 0 | . | . | . | . |

**Type 3 Tests of Fixed Effects**

| Effect | Num DF | Den DF | F Value | Pr > F |
|---|---|---|---|---|
| Treatment | 2 | 25.1 | 26.86 | <.0001 |
| Time | 4 | 25 | 1.15 | 0.3562 |

**Least Squares Means**

| Effect | Treatment | Estimate | Standard Error | DF | t Value | Pr > \|t\| |
|---|---|---|---|---|---|---|
| Treatment | dsRNA + DNA | 6692.60 | 1713.45 | 2.49 | 3.91 | 0.0414 |
| Treatment | dsRNA+saliva | 2953.07 | 1713.45 | 2.49 | 1.72 | 0.2017 |
| Treatment | dsRNA+water | 9831.64 | 1712.91 | 2.48 | 5.74 | 0.0174 |

**Differences of Least Squares Means**

| Effect | Treatment | Treatment | Estimate | Standard Error | DF | t Value | Pr > \|t\| | Adjustment | Adj P |
|---|---|---|---|---|---|---|---|---|---|
| Treatment | dsRNA + DNA | dsRNA+saliva | 3739.53 | 822.47 | 25 | 4.55 | 0.0001 | Tukey-Kramer | 0.0003 |
| Treatment | dsRNA + DNA | dsRNA+water | -3139.04 | 963.43 | 25.2 | -3.26 | 0.0032 | Tukey-Kramer | 0.0087 |
| Treatment | dsRNA+saliva | dsRNA+water | -6878.57 | 963.43 | 25.2 | -7.14 | <.0001 | Tukey-Kramer | <.0001 |

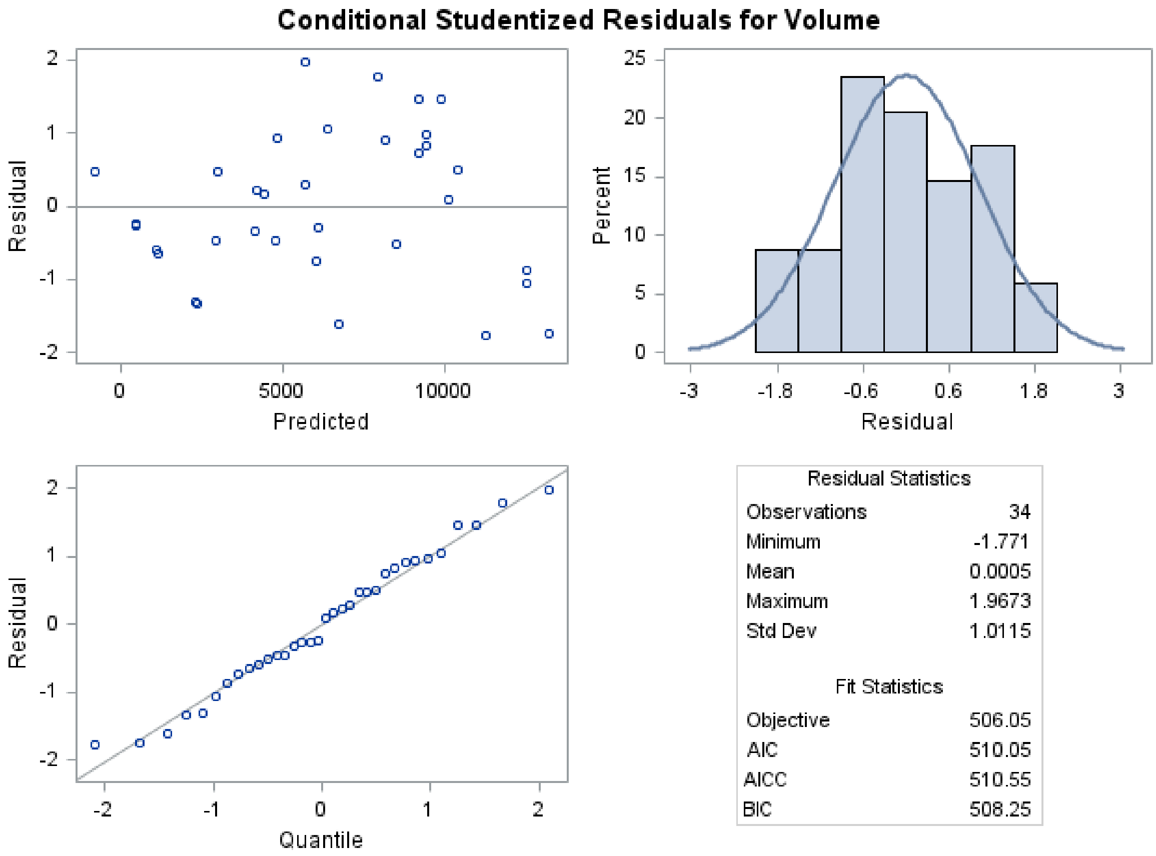

Residual Statistics

| | |
|---|---|
| Observations | 34 |
| Minimum | -1.771 |
| Mean | 0.0005 |
| Maximum | 1.9673 |
| Std Dev | 1.0115 |

Fit Statistics

| | |
|---|---|
| Objective | 506.05 |
| AIC | 510.05 |
| AICC | 510.55 |
| BIC | 508.25 |

### The SAS System

#### The Mixed Procedure

| Model Information | |
|---|---|
| Data Set | WORK.GEL_DATA |
| Dependent Variable | Volume |
| Covariance Structure | Variance Components |
| Estimation Method | REML |
| Residual Variance Method | Profile |
| Fixed Effects SE Method | Kenward-Roger |
| Degrees of Freedom Method | Kenward-Roger |

| Class Level Information | | |
|---|---|---|
| Class | Levels | Values |
| Treatment | 2 | dsRNA + SG dsRNA+water |
| Time | 5 | 0 1 10 30 60 |
| Gels | 5 | 4 5 6 7 8 |
| Life_stage | 5 | 2 3 4 5 Adult |
| Repetition | 1 | A |

| Dimensions | |
|---|---|
| Covariance Parameters | 2 |
| Columns in X | 8 |
| Columns in Z | 5 |
| Subjects | 1 |
| Max Obs per Subject | 88 |

| Number of Observations | |
|---|---|
| Number of Observations Read | 90 |
| Number of Observations Used | 88 |
| Number of Observations Not Used | 2 |

| Iteration History | | | |
|---|---|---|---|
| Iteration | Evaluations | -2 Res Log Like | Criterion |
| 0 | 1 | 1563.75983248 | |
| 1 | 3 | 1533.00315077 | 0.00003938 |
| 2 | 1 | 1532.97271701 | 0.00000169 |
| 3 | 1 | 1532.97151484 | 0.00000000 |

Convergence criteria met.

| Covariance Parameter Estimates |
|---|

| Cov Parm | Estimate |
|---|---|
| Gels | 4616109 |
| Residual | 5537784 |

| Fit Statistics | |
|---|---|
| -2 Res Log Likelihood | 1533.0 |
| AIC (Smaller is Better) | 1537.0 |
| AICC (Smaller is Better) | 1537.1 |
| BIC (Smaller is Better) | 1536.2 |

**Solution for Fixed Effects**

| Effect | Treatment | Time | Estimate | Standard Error | DF | t Value | Pr > \|t\| |
|---|---|---|---|---|---|---|---|
| Intercept | | | 10922 | 1141.51 | 6.54 | 9.57 | <.0001 |
| Treatment | dsRNA + SG | | -11818 | 530.11 | 78 | -22.29 | <.0001 |
| Treatment | dsRNA+water | | 0 | . | . | . | . |
| Time | | 0 | 132.44 | 981.55 | 78 | 0.13 | 0.8930 |
| Time | | 1 | 546.51 | 755.19 | 78.1 | 0.72 | 0.4714 |
| Time | | 10 | 583.59 | 744.16 | 78 | 0.78 | 0.4353 |
| Time | | 30 | 159.96 | 744.16 | 78 | 0.21 | 0.8304 |
| Time | | 60 | 0 | . | . | . | . |

**Type 3 Tests of Fixed Effects**

| Effect | Num DF | Den DF | F Value | Pr > F |
|---|---|---|---|---|
| Treatment | 1 | 78 | 496.95 | <.0001 |
| Time | 4 | 78 | 0.23 | 0.9189 |

**Least Squares Means**

| Effect | Treatment | Estimate | Standard Error | DF | t Value | Pr > \|t\| |
|---|---|---|---|---|---|---|
| Treatment | dsRNA + SG | -611.22 | 1058.49 | 4.86 | -0.58 | 0.5894 |
| Treatment | dsRNA+water | 11206 | 1034.96 | 4.44 | 10.83 | 0.0002 |

**Differences of Least Squares Means**

| Effect | Treatment | Treatment | Estimate | Standard Error | DF | t Value | Pr > \|t\| | Adjustment | Adj P |
|---|---|---|---|---|---|---|---|---|---|
| Treatment | dsRNA + SG | dsRNA+water | -11818 | 530.11 | 78 | -22.29 | <.0001 | Tukey-Kramer | <.0001 |

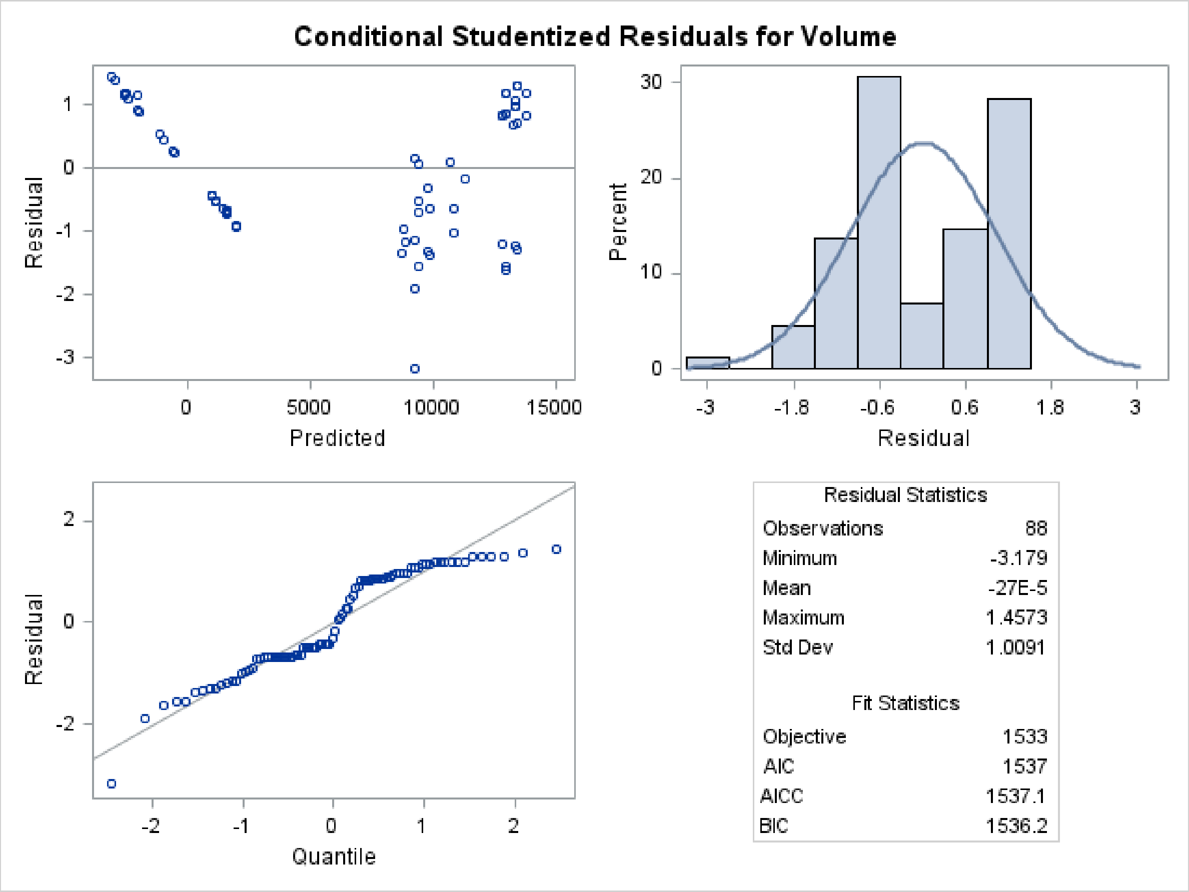
Conditional Studentized Residuals for Volume
Residual
1
0
-1
-2
-3
0
5000
10000
15000
Predicted
Percent
30
20
10
0
-3
-1.8
-0.6
0.6
1.8
3
Residual
Residual
2
0
-2
-2
-1
0
1
2
Quantile
Residual Statistics
Observations 88
Minimum -3.179
Mean -27E-5
Maximum 1.4573
Std Dev 1.0091
Fit Statistics
Objective 1533
AIC 1537
AICC 1537.1
BIC 1536.2
